# Human RNase H2 upregulation counteracts oncogene- and chemotherapy-induced replication stress

**DOI:** 10.1101/2024.12.16.628316

**Authors:** Rosanna J. Wilkins, Abirami Kannan, Siobhan A. Plass, Claire Wilson, Richard D. W. Kelly, Claire H. M. Tang, Panagiotis Kotsantis, Martin A. M. Reijns, Aditi Kanhere, Eva Petermann

**Affiliations:** Department of Cancer and Genomic Sciences, College of Medicine and Health, University of Birmingham, UK; Birmingham Centre for Genome Biology, University of Birmingham, UK; Department of Molecular & Clinical Cancer Medicine, Institute of Systems, Molecular & Integrative Biology, University of Liverpool, UK; MRC Human Genetics Unit, Institute of Genetics and Cancer, University of Edinburgh, UK; Division of Biomedical and Life Sciences, Faculty of Health and Medicine, Lancaster University, UK

## Abstract

RNase H2 is a heterotrimeric endoribonuclease that resolves RNA:DNA hybrids and genome-embedded ribonucleotides, which are implicated in DNA replication stress and cancer development. Protein and/or mRNA levels of individual RNase H2 subunits are elevated in some cancers, but little is known about the mechanisms or consequences of RNase H2 upregulation. We report that RNase H2 subunits are upregulated at the protein level in response to replication stress induced by oncogenes and chemotherapy drugs in human cancer and non-cancer cell lines. We show that inducible overexpression of the RNASEH2B subunit increases levels of the active RNase H2 heterotrimer. While causing only subtle changes to gene expression, RNASEH2B overexpression is unexpectedly associated with increased RNA:DNA hybrid levels. RNASEH2B overexpression prevents further increases in RNA:DNA hybrid levels by camptothecin or hydroxyurea and reduces replication fork stalling in presence of these drugs. Surprisingly, RNase H2 levels do not strongly impact survival of chemotherapy treatments but appear to have more subtle effects on genome instability and innate immune signalling. In contrast, increased RNase H2 levels in presence of oncogenic HRAS limit not only RAS-induced replication fork stalling but also cell death. Our findings shed new light on the functions of RNase H2 and suggest that upregulation of RNase H2 may be an important aspect of replication stress responses in cancer.

## Introduction

Replication stress can contribute to tumorigenesis via genome instability^1^ but is also a therapeutic target and a central mechanism of action for many cytotoxic cancer therapies^2^. Nuclear processes such as transcription can contribute to replication stress by generating barriers to replication fork progression. Transcription can interfere with replication fork progression through topological stress, collisions of the RNA and DNA polymerase machineries or co-transcriptional RNA:DNA hybrids called R-loops^3^.

Oncogenes such as HRAS, SS18-SSX1 and EWS–FLI1 increase RNA:DNA hybrid levels and transcription-replication conflicts (TRCs)^4–6^. Chemotherapies can also induce RNA:DNA hybrids. For instance, camptothecins (CPT) inhibit DNA topoisomerase I (TOP1), which alleviates supercoiling, increasing the likelihood of R-loop formation^7^. Gemcitabine and hydroxyurea deplete deoxyribonucleotides which can lead to increased ribonucleotide incorporation^8,9^ and stalled replication forks, which can in turn promote R-loop formation^10^. RNA:DNA hybrids can thereby contribute to DNA damage, genome instability and cell death caused by chemotherapies. They may also contribute to the toxicity of targeted cancer therapies such as ATR inhibitors^11^ and to innate immune activation^12^.

R-loop homeostasis is a fine balance between R-loop formation and removal. In eukaryotes, RNaseH1 and RNase H2 can digest the RNA component of RNA:DNA hybrids^13^. RNase H2 is a heterotrimer where RNASEH2A contains the catalytic domain, whereas RNASEH2B and RNASEH2C are essential accessory subunits^14,15^. RNASEH2B is required for the nuclear localisation of the RNase H2 complex and interacts with the DNA sliding clamp, PCNA^16,17^. RNase H2 initiates ribonucleotide excision repair (RER), the removal of mis-incorporated single ribonucleotides from duplex DNA^18^, and can remove R-loops (Fig. 1A). RNase H2 is the predominant RNase H activity in yeast and mammals^19,20^. In *S. cerevisiae* RNase H2 protein levels are cell-cycle regulated^21,22^ and R-loops at replication forks are thought to be processed specifically by RNase H2^23^. However, in mammals RNase H2 levels appear to be constant throughout the cell cycle^8,9^ and the contribution of this enzyme to R-loop removal is less well studied. RNase H2 deficiency can cause neuroinflammation^24^ or contribute to cancer development^25,26^, whereas RNase H2 upregulation is associated with cancer progression^27–29^. R-loop interacting proteins such as RNase H2 are potential therapeutic targets in cancer^29^.

**Figure 1.**
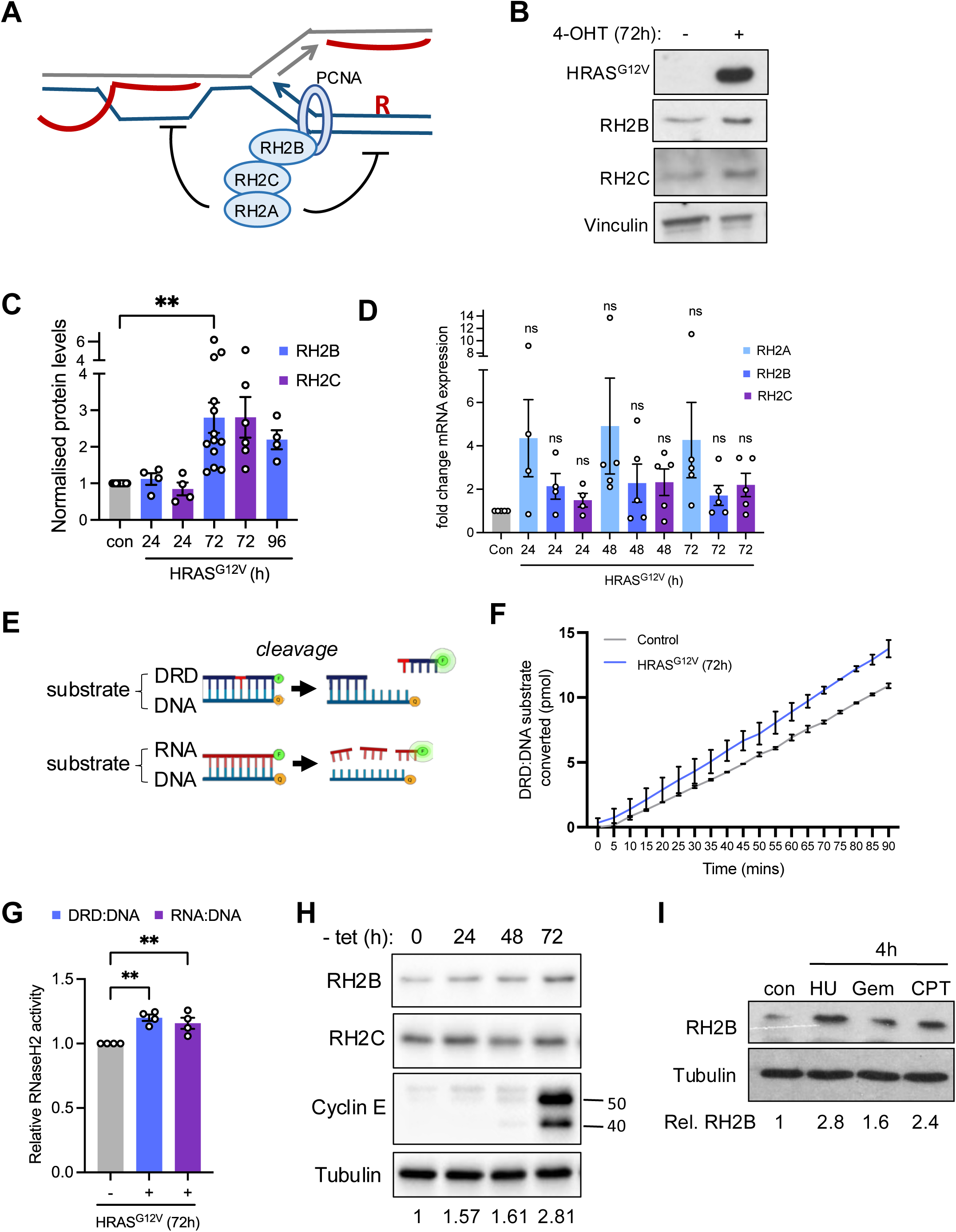
Oncogene-induced replication stress increases RNase H2 protein levels and -activity. (A) RNase H2 complex subunits and functions. (B) Protein levels of HRAS, RNASEH2B (RH2B), RNASEH2C (RH2C) and Vinculin (loading control) in BJ-hTERT HRAS^V12ERTAM^ cells +/-72 h HRAS^G12V^ induction with 4-hydroxytamoxifen (4-OHT). (C) Relative protein levels of RH2B and RH2C after 24, 72 or 96 h HRAS^G12V^ induction normalised to ethanol (con) for each protein. N=5 (24 and 96 h), N=13 (RH2B 72 h), N=6 (RH2C 72 h). (D) RT-qPCR analysis of *RNASEH2A, RNASEH2B*, and *RNASEH2C* expression after HRAS^G12V^ induction. Asterisks compare to con. N = 4 (24 h), N = 5 (48 and 72 h). (E) Substrates used for the RNase H2 activity assay with forward strand covalently coupled to 3’-fluorescein (green) and reverse strand linked to 5’-DABCYL quencher (orange). (F) Representative time course of DRD:DNA substrate conversion during incubation with whole cell extract +/- 72 h HRAS^G12V^ induction (control). N = 2 measured on the same plate. (G) Relative RNase H2 activities in cell extracts +/- 72 h HRAS^G12V^ induction. N = 4. (H) Protein levels of CYCLIN E, RH2B, RH2C and Tubulin (loading control) in U2OS-CYCLIN E cells +/-CYCLIN E induction with tetracycline removal (-tet) for the times indicated. (I) Protein levels of RH2B and Tubulin (loading control) in BJ-hTERT HRAS^V12ERTAM^ cells without HRAS^G12V^ induction after 4 h treatment with 200 μM hydroxyurea (HU), 25 nM gemcitabine (GEM), 10 μM camptothecin (CPT) or DMSO (con). Means and SEM (bars) of independent experiments are shown. Asterisks indicate p-values (ANOVA or mixed-effects analysis, * p < 0.05, ** p < 0.01).

RNase H2 deficiency has long been studied due to its association with the neuroinflammatory disorder Aicardi-Goutières Syndrome^15,17^. RNase H2 loss causes spontaneous replication stress^15^, and interferes with the proper processing and progression of replication forks in presence of endogenous replication stress-inducing agents^30,31^. However, RNase H2 protein levels are often increased in cancers such as colon cancer^29,32^ where they can contribute to aggressive phenotypes such as cell migration and growth^33,34^. The roles of increased RNase H2 activity during replication stress responses are not well understood.

Here we investigate roles of RNase H2 in response to replication stress induced by oncogene activation and chemotherapy drugs, both in cancer and non-cancer cells. We show that RNase H2 protein levels and activity are up-regulated in response to replication stress. Using inducible non-cancer and cancer cell models, we show that overexpression of the RNASEH2B subunit can increase protein levels and activity of the entire RNase H2 complex. RNASEH2B overexpression unexpectedly increases global RNA:DNA hybrid levels, but prevents further RNA:DNA hybrid increases after treatment with chemotherapy drugs. RNASEH2B overexpression counteracts drug- and oncogene-induced replication fork stalling, with potential implications for genome instability, cytosolic nucleic acid sensing, and cell survival.

## Materials and Methods

### Cell lines and reagents

U2OS cells were obtained from ATCC. HCT116, HT55, NCIH747 and LS174T cells were from the Charles Swanton lab^1^. Human BJ-hTERT HRASV12^ER-TAM^ were from the Agami and de Vita labs^35^. U2OS-Cyclin E cells were from the Jiri Lukas lab^36^. Cell lines were authenticated using 17-locus STR profiling (LGC Standards) and verified to be Mycoplasma-free by PCR. All cell lines were grown in Dulbecco’s modified Eagle’s Medium (DMEM), supplemented with 10% foetal bovine serum and 1% L-glutamine, in a humidified atmosphere containing 5% CO2. U2OS-Cyclin E cells were also grown in presence of G418 (400 µg/ml), puromycin (1 µg/ml), and tetracycline (2 µg/ml). HRAS^V12^ was induced with 500 nM 4-hydroxytamoxifen (4OHT, Sigma). RNASEH2A, RNASEH2B or RNASEH2C were induced with 1 µg/ml doxycycline (Sigma). Cyclin E was induced by washing cells three times in PBS and transferring into medium containing 10% tetracycline-free fetal calf serum (PAA-GE Healthcare), G418 and puromycin. Camptothecin, hydroxyurea and 5,6-dichloro-1-β-D-ribofuranosylbenzimidazole (DRB) were from Sigma. Gemcitabine and triptolide were from Tocris Bioscience.

### Lentiviral transfections

Lentiviral vectors were generated by Genscript Biotech, using pLIX403-ccdB-Blast, a gift from Alejandro Chavez (Addgene plasmid # 158560; http://n2t.net/addgene:158560; RRID:Addgene 158560), to generate pLIX403-ccdB-Blast-RNASEH2B. 10 µg purified vector DNA, lentiviral packaging plasmids (RRE, REV and VSV) and lipofectamine 2000 (Invitrogen) were incubated for 20 min at room temperature in OptiMEM (Gibco) before being added dropwise to HEK293FT cells in antibiotic-free DMEM and incubated for 48 h. Collected medium was passed through a 0.45 µm filter and supplemented with 8 µg/ml polybrene (Sigma), then mixed with cells followed by centrifugation for 90 min at 4000 rpm (3739 rcf), before replacing with fresh medium and repeating the polybrene infection 24 h later. Cells were washed with PBS and selected with 10 µg/ml blasticidin (Sigma) for 7 days. Clones were isolated by plating selected cells at 1000 cells per 10 cm dish.

### siRNA transfections

Cells were seeded in 6-well plates and incubated with DharmaFECT 1 reagent (Horizon Discovery) and 50 nM siRNA per well. ‘All stars negative control siRNA’ was purchased from Qiagen. ON-TARGETplus SMARTpool siRNAs against RNASEH2A (L-003535-01-0005), RNASEH2B (L-014369-01-0005) and RNASEH2C (L-014801-00-0005) were purchased from Horizon Discovery.

### Western blot

For RNASEH2B detection, cells should not be freeze-thawed prior to extraction to avoid loss of protein. Cells were extracted in UTB buffer (50mM Tris-HCl pH 7.5, 150mM β-mercaptoethanol and 8 M urea) unless indicated otherwise, sonicated and protein concentration determined by Bradford assay. Primary antibodies were: Rabbit anti-RNASEH2B (Invitrogen PA5-59095, 1:250; Abcam ab122619, 1:250; Invitrogen PA5-83560, 1:250), anti-RNASEH2C (Proteintech 16518-1-AP, 1:500), rabbit anti-RNASEH2A (Proteintech 16132-1-AP, 1:2000), mouse anti-HRAS (Santa Cruz sc-29, 1:1,000), rabbit anti-MYC-tag (Cell Signaling 2272, 1:1,000), mouse anti-αtubulin (B512 Sigma T6074, 1:40000), rabbit anti-vinculin (Abcam ABCAAB1 29002-100, 1:10,000), rabbit anti-TBK1 (Cell Signaling 3504S, 1:5,000), rabbit anti-phospho(S172) TBK1 (Cell Signaling 5483S, 1:5,000). Secondary antibodies were goat anti-rabbit-HRP (Cell Signaling, 7074, 1:5,000), goat anti-mouse-HRP (Cell Signaling 7076, 1:5,000). Protein quantification of scanned blots was performed using ImageJ (http://rsbweb.nih.gov/ij/)^37^.

### RNase H2 activity assay

Desalted oligonucleotides were from Eurogentec (Liege, Belgium) and were reconstituted at 100 µM in sterile nuclease-free water. Forward strands: DRD, 5’-GATCTGAGCCTGGGaGCT-3’, DNA, 5’-GATCTGAGCCTGGGAGCT-3’, RNA, 5’-gaucugagccugggagcu-3’. Reverse strand: DNA, 5’-AGCTCCCAGGCTCAGATC-3’. Lower case denotes RNA. Forward strands have a 3’-fluorescein residue and reverse strands have a 5’ DABCYL quencher residue. Oligonucleotides were annealed by denaturing 10 µM forward strand and 10 µM reverse strand at 95 °C for 5 min in 1X oligo buffer (60mM KCl, 50mM Tris-HCl pH 8) then gradually cooling them to room temperature. Standard curves comprise forwards strands being annealed to reverse strands that lack the 5’-DABCYL quencher. Cells were washed twice with 1X PBS, trypsinized, and pelleted at 845 g for 1 minute at 4 °C. Cell pellets were snap frozen in dry ice and stored at -80°C. Cell extracts were prepared using whole cell extract buffer (50mM Tris pH8, 280mM NaCl, 0.5% NP40, 0.2mM EDTA, 0.2mM EGTA and 10% glycerol) for 10 min on ice followed by adding an equal volume of CB buffer (20mM HEPES pH7.9, 10mM KCl, 1mM EDTA and 10% glycerol) for 10 min on ice. Both lysis buffers were supplemented with 1mM phenylmethylsulfonyl fluoride, 1mM dithiothreitol, 1X protease inhibitor cocktail (Roche) and 1X phosSTOP (Roche) prior to use. Cell lysates were centrifuged for 10 min at 21130 g and 4 °C and protein concentration measured by Bradford assay. Lysates were diluted in RNase H2 assay (RA) buffer (50mM Tris pH8, 60mM KCl, 10mM MgCl2, 0.01% BSA and 0.01% Triton X-100) to a final protein concentration of 100 ng/µl and incubated with 250 nM substrate in black 96-well flat-bottomed plates (Costar) in technical triplicates. Fluorescence intensity was measured (10 flashes per well), using 485 nm excitation and 520 nm emission filters, every 5 min for 90 min at 24°C on a BMG Labtech PHERAstar FS multimode plate reader or a Perkin-Elmer EnSpire multimode plate reader. Standard curves were generated to convert relative fluorescence units (RFU) to pmol of substrate converted by linear regression. For absolute values, correction was performed by subtracting the negative control wells, containing substrate only. For the DRD:DNA substrate, values from DNA:DNA substrate wells were subtracted for additional correction.

### Co-immunoprecipitation

Immunoprecipitation was carried out using the ChromoTek Myc-trap® Magnetic Agarose Kit (Proteintech) according to the manufacturer’s instructions with some modifications. Buffers provided in the kit were supplemented with DNase I, 2.5 mM MgCl2, 1X EDTA-free protease inhibitor cocktail (Roche), 1X phosSTOP (Roche) and 1 mM phenylmethylsulfonyl fluoride (ThermoFisher). A total of 1 mg protein per condition was incubated with the anti-Myc-tag nanobody for 1 h. Supernatants were eluted in 5X Laemmli buffer and denatured at 95 °C for 5 min. Samples were analysed with SDS-PAGE and Western blot.

### Slot blot

Genomic DNA was extracted using the DNeasy Blood & Tissue kit (Qiagen) according to the manufacturer’s instructions. 10 µg genomic DNA per sample was mock treated or treated with 2 U/µg DNA of RNaseH (NEB, M0297) for 2 h at 37 °C. 250 ng DNA per well was loaded onto the slot blot apparatus, and transferred onto a pre-wetted nylon membrane (Amersham Hybond N+) by vacuum suction. The membrane was UV-crosslinked at 1200 µJ and blocked in sterile 5 % milk/TBST for 1 h at room temperature. The membrane was incubated overnight at 4°C in mouse anti-DNA-RNA hybrid antibody (S9.6 hybridoma growth medium, ATCC HB-8730, 1:1,000) or mouse anti-dsDNA antibody (Abcam ab27156, 1:100,000) in sterile 5 % BSA/TBST, before being washed in TBST and incubated in goat anti-mouse HRP (Cell Signaling 7074, 1:5000) 5 % milk/TBST for 1 h at room temperature. Images were taken using a ChemiDoc™ MP Imaging System (BioRad) and quantifications performed using ImageJ.

### DNA fibre assay

Cells were pulse-labelled with 25 µM CldU followed by 250 µM IdU as indicated. DNA fibres were spread by spotting 2 µl of cells at 5 x 10^5^ cells/ml in PBS onto microscope slides (Menzel Gläser) and allowed to settle for 5 min. Cells were mixed with 7 µl of spreading buffer (0.5% SDS, 200 mM Tris-HCl pH 7.4 and 50 mM EDTA) for 2 min. DNA fibres were spread by tilting the slides for 3 min and fixed in methanol/ glacial acetic acid (3:1). Rehydrated fibre spreads were denatured with 2.5 M HCl for 75 min at room temperature. Slides were blocked with 1% BSA and 0.1% Tween-20 in PBS before incubation with rat anti-BrdU (BU1/75, Abcam ab6326, 1:700) and mouse anti-BrdU (B44, Becton Dickinson 347580, 1:500) for 1 h followed by fixation with 4% PFA for 10 min. Slides were incubated with secondary antibodies anti-mouse IgG AlexaFluor 488 (ThermoFisher, 1:500) and anti-rat IgG AlexaFluor 594 (Invitrogen, 1:500). Slides were mounted (Fluoroshield, Sigma) and imaged using a Nikon E600 microscope with a Nikon Plan Apo 60x (1.3NA) oil lens, an Andor Zyla sCMOS digital camera and the Nikon NIS elements BR 5.41.01 software. The lengths of the CldU and IdU tracts were measured using the ImageJ software and lengths converted into μm using a micrometer slide. For replication structures, stalled forks were quantified by scoring the percentages of CldU-only labelled fibres of other quantifiable structures.

### Immunofluorescence

Cells were washed and fixed to coverslips with 4% paraformaldehyde for 10 min at room temperature, then permeabilized with 0.25% Triton X-100 in PBS for 5 min at room temperature before blocking with 1% bovine serum albumin and 0.1% Tween-20 in PBS. For ssDNA and dsDNA staining, coverslips were fixed and permeabilized, then treated with 0.1 mg/ml RNase A (ThermoFisher, EN0531) for 30 min at 37 °C and 5% CO2. Primary antibodies were mouse anti-ssDNA (Merck MAB3868, 1:1,000), mouse anti-dsDNA (Merck MAB1293, 1:100), rabbit anti-53BP1 (Bethyl A300-272A, 1:3000), mouse anti-phospho-Histone H2AX (Merck JBW301, 1:1000), rabbit anti-RAD51 (Abcam ab63801, 1:800). Secondary antibodies were anti-mouse IgG AlexaFluor 488 (ThermoFisher, 1:500) and anti-rabbit IgG AlexaFluor 594 (ThermoFisher, 1:500). Coverslips were mounted in Fluoreshield with 4.6-diamidino-2-phenylindole (DAPI) (Sigma). For quantification of γH2AX nuclear intensity, nuclear masks were created using ImageJ software based on DAPI-staining and mean fluorescence intensities per pixel were quantified per nucleus.

### RNA extraction

Cells were harvested in TRI Reagent® (Zymo Research) and RNA extraction was performed using the Direct-zol RNA Miniprep Kit (Zymo Research), including DNase I treatment, according to the manufacturer’s instructions. RNA quality and concentration was determined using the 4200 tapestation system.

### Reverse Transcription Quantitative real-time PCR (RT-qPCR)

Total RNA (1 ug) was reverse-transcribed using qScript cDNA synthesis kit (Quantabio), following the manufacturer’s instructions. RT-qPCR primers for amplification were obtained from Merck and their sequences are as follows: *ACTB* Forward TTGCGTTACACCCTTTCTTG, *ACTB* Reverse CACCTTCACCGTTCCAGTTT;

RNASEH2A Forward AATGGAGGACACGGACTTTG, *RNASEH2A* Reverse ATGTCTCTGGCATCCCTACG;

RNASEH2B Forward CTGGTGACCAAGCTTCCACT, *RNASEH2B* Reverse ATGGAGGATTTGGCAATGAG;

*RNASEH2C* Forward CTTAACTTGGCCCAGCCTTG, *RNASEH2C* Reverse TCCTTTATTGGGGTGATGGA.

2 ul of cDNA was incubated with primers and SYBR™ Green PCR Master Mix (ThermoFisher) and analysed using the Real Time PCR QuantStudio 5 system (ThermoFisher). Cycling parameters were 50 °C for 2 min and 95 °C for 5 min followed by 40 cycles of 95 °C for 30 seconds, 60 °C for 30 seconds and 72 °C for 30 seconds. Results were normalized to *ACTB*.

### Steady-state eukaryotic directional mRNA-sequencing

Library preparation and sequencing on the Illumina platform NovaSeq 6000 S4 (PE150) were performed by Novogene. Sequences were first quality checked using FastQC v0.11.9 (https://www.bioinformatics.babraham.ac.uk/projects/fastqc/), then quality filtered and trimmed using TrimGalore v0.6.6 (https://www.bioinformatics.babraham.ac.uk/projects/trim_galore/). Genome mapping was performed with STAR^38^ (v2.7.10b) against the human reference genome (hg38 assembly). Reads were counted using FeatureCounts^39^ (subread v2.0.3) and differential analysis was performed using DESeq2^40^, with p-value calculated using the Wald test. Differential genes were identified by P-value<0.01 and LogFC>2. Volcano plots were generated using EnhancedVolcano.

Normalised read counts were converted into GCT files compatible with GSEA 4.3.3 (Broad Institute)^41,42^. GSEA was run with the following parameters: number of permutations (1000), permutation type (gene_set), ChIP platform (Human_Ensembl_Gene_ID_MSigDB.v2024.1.Hs.chip). All other parameters were left as the default setting. The gene sets databases used were from the Molecular Signatures Database (MSigDB)^43,44^.

### Colony survival assay

For siRNA transfections, HCT116 cells at 70% confluency were transfected for 24 h prior to being plated. Cells were allowed 24 h to adhere before doxycycline or drug treatments. Doxycycline was replenished every 2 days. To stain colonies, cells were washed twice with 1X PBS and fixed in 50% ethanol, 2% methylene blue for 20 min at room temperature. Colonies of >50 cells were counted. For cell proliferation assays, 20,000 cells were seeded per well of a 6-well plate. After treatments, cells were trypsinized, resuspended in 1ml of PBS and counted using cell counting chambers.

### Flow cytometry

Cells were fixed in 70% ice-cold ethanol and centrifuged at 250 g for 5 min. For inclusion of the sub-G1 population, culture medium and pre-trypsinisation washes were also collected. Cells were washed 3 times with PBS prior to incubating with 10 µg/ml propidium iodide and 25 µg/ml ribonuclease A from bovine pancreas (Merck) in PBS. Cell cycle profiles were generated using the Fortessa X-20 flow cytometer and analysed with the BD FacsDiva software.

### TCGA data analysis

Cancer Genome Atlas datasets were accessed using cBioPortal. Colorectal Adenocarcinoma (COAD, TCGA, PanCancer Atlas), all complete samples, was interrogated for *RNASEH2A*, *RNASEH2B* and *RNASEH2C* mRNA expression z-scores relative to diploid samples (RNA Seq V2 RSEM). The correlation with aneuploidy scores was obtained from the built-in cBioPortal Comparison/Survival tab.

### Statistical analysis

Unless specified, all values for bar graphs are the mean +/- s.e.m of results from biological replicates. Scatter plots represent individual data points and medians from independent biological repeats. Numbers of independent biological repeats (N) are indicated in the figure legends. Statistical tests were performed using the GraphPad Prism 10 software, version 10.2.0 (392). Test were chosen depending on the structure of the datasets, using a paired or unpaired student’s *t*-test for two comparisons or a one-or two-way ANOVA or mixed-effects analysis with Dunnett’s or Tukey’s for multiple comparisons. For analysis of TCGA datasets, the cBioPortal in-built Wilcoxon Test was used.

## Results

### Induction of RNase H2 protein levels and activity in response to oncogene-induced replication stress

We had previously shown that the expression of oncogenic HRAS^G12V^ in BJ-hTERT fibroblasts increased RNA:DNA hybrid levels, caused TRCs and increased mRNA and protein levels of RNaseH1^6^. We therefore decided to further investigate how HRAS^G12V^ induction influences the protein levels of RNA:DNA hybrid-resolving enzymes. We found that HRAS^G12V^ induction for 3 or 4 days, but not for 24 h, increased protein levels of RNASEH2B and RNASEH2C (Fig. 1B, C; Fig. S1A, B). HRAS^G12V^ induction was accompanied by a trend towards increased mRNA expression of genes encoding RNase H2 subunits, especially of *RNASEH2A* (Fig. 1D).

To measure the impact of oncogenic HRAS^G12V^ on cellular RNase H2 activity, we employed a fluorescence resonance energy transfer (FRET)-based activity assay^14^ to measure nuclease activity against RNA:DNA hybrid substrate or double stranded DNA with a single ribonucleotide (DRD:DNA) (Fig. 1E). As RNase H2 is the only known enzyme to specifically cleave DNA-embedded ribonucleotides, the DRD:DNA substrate is considered a specific readout of RNase H2 activity^8^. Both RNase H2-specific and RNA:DNA hybrid resolution activities were increased after HRAS^G12V^ induction (Fig. 1F, G).

HRAS^G12V^ will accelerate S phase entry, which may increase RNase H2 protein levels due to cell cycle regulation. However, the biggest increase in S/G2 phase percentage occurred 24 h after HRAS^G12V^ induction, when RNASEH2B protein levels were not elevated (Fig. S1C, D). We also previously showed that MAP kinase signalling and replication stress are much more strongly elevated after 72 h compared to 24 h HRAS^G12V^ induction ^6^. Therefore, RNase H2 protein levels increase in response to RAS signalling and/or replication stress rather than S phase entry.

To further investigate whether the increase in RNase H2 protein and activity was specific to RAS signalling or more generally observed during oncogene-induced replication stress, we used U2OS cells inducibly overexpressing the oncogene Cyclin E ^45^, which also increased RNASEH2B protein levels (Fig. 1H). We next used short 4 h treatments with replication stress-inducing drugs hydroxyurea (HU), gemcitabine (GEM), or camptothecin (CPT) in BJ-hTERT cells without HRAS^G12V^ induction. This also increased RNASEH2B protein levels (Fig. 1I). Taken together, these data suggest that RNase H2 subunits are up-regulated at the protein level in response to replication stress.

### Induction of RNase H2 protein and activity in response to drug-induced replication stress

We further investigated the role of RNase H2 in response to replication stress-inducing drugs using colon cancer (CRC) cell lines relevant to chemotherapy drug responses and cancers with elevated RNase H2 subunit levels^29,32^. mRNA expression of at least one RNase H2 subunit, most commonly *RNASEH2B*, is increased in over 25% of CRCs which correlates with higher aneuploidy, and therefore likely with higher chromosomal instability and possibly higher replication stress^1^ (Fig. 2A, B). We used four CRC cell lines. HT55 and NCIH747 are characterised by high chromosomal instability and replication stress, while LS174T and HCT116 cells are characterised by microsatellite instability and low replication stress^1^. We then investigated protein levels of RNASEH2B and RNASEH2C after 2 h treatment with HU, GEM or CPT (Fig. 2C-J; Fig. S2A-H). Despite relatively high variability between experiments, there was an overall trend of drug-induced increases of RNASEH2B and RNASEH2C levels across most cell lines. Although 2 h CPT treatment increased RNASEH2B protein levels in HCT116 cells (Fig. 2I), in contrast to HRAS activation this was not accompanied by a trend towards increased *RNASEH2A* mRNA expression (Fig. 2K). RNase H2 activity increased in response to replication stress inducing agents in selected cell lines, HT55 and HCT116, in line with the increases in protein levels (Fig. 2L, M).

**Figure 2.**
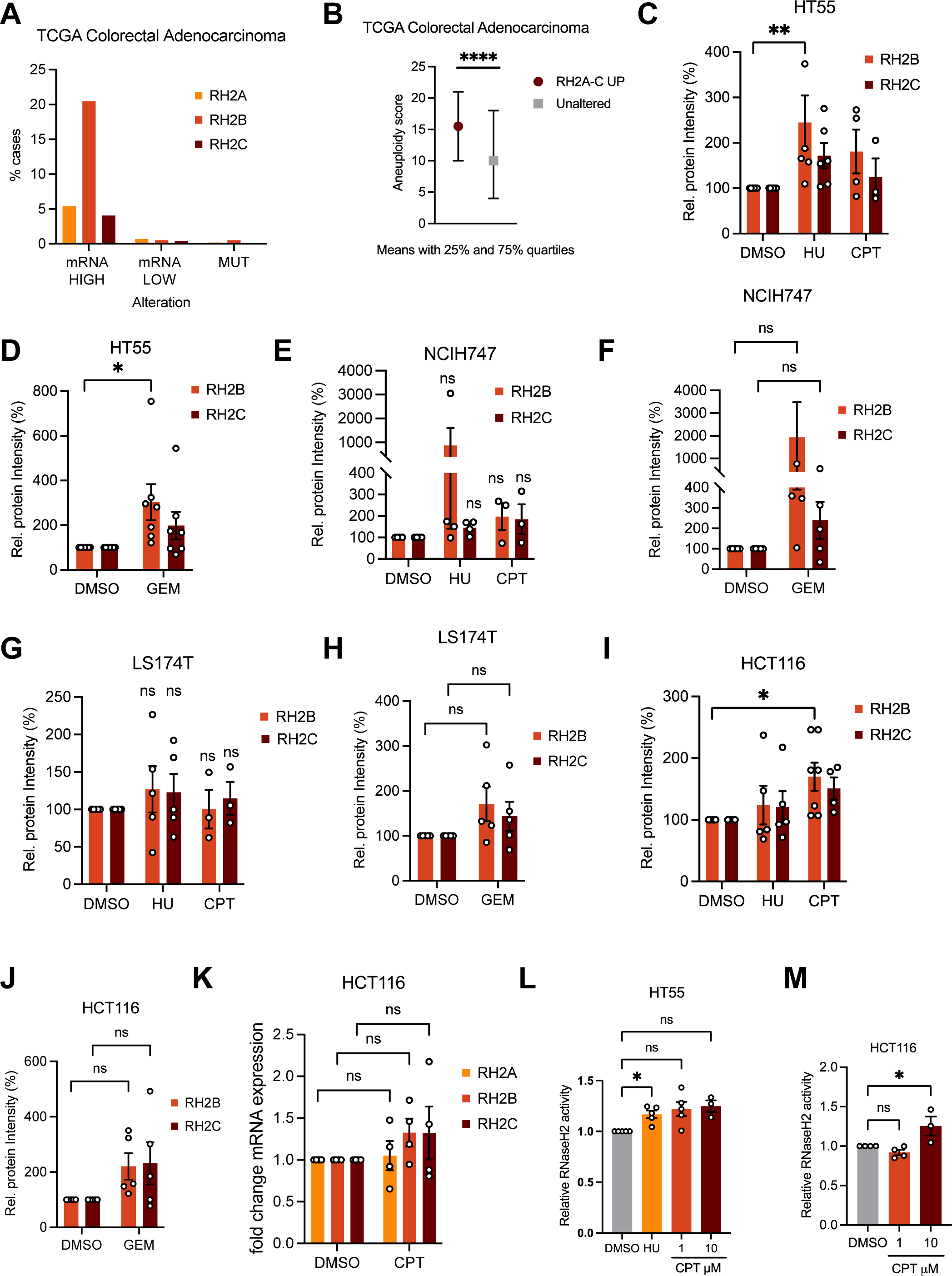
RNase H2 protein levels and -activity increase in response to chemotherapeutic drugs. (HA) *RNASEH2A, -B, and -C* mRNA levels are increased in colorectal adenocarcinoma (TCGA Pan-Cancer Atlas^68,69^, RNA Seq V2 RSEM, z-score threshold ± 2.0). (B) Increased *RNASEH2A, -B, and -C* mRNA levels are correlated with higher aneuploidy in colorectal adenocarcinoma (TCGA Pan-Cancer Atlas). Medians with 25% and 75% quartiles are shown. Wilcoxon Test, **** p < 0.0001. (C) Relative protein levels of RH2B and RH2C in HT55 cells after 2 h treatment with 200 μM HU, 10 μM CPT or DMSO. N = 3-7. (D) Relative protein levels of RH2B and RH2C in HT55 cells after 2 h treatment with 25 nM GEM or DMSO. N = 7. (E) Relative protein levels of RH2B and RH2C in NCIH747 cells after treatment as in C. N = 3-4. (F) Relative protein levels of RH2B and RH2C in NCIH747 cells after treatment as in D. N = 5. (G) Relative protein levels of RH2B and RH2C in LS174T cells after treatment as in C. N = 3-5. (H) Relative protein levels of RH2B and RH2C in LS174T cells after treatment as in D. N = 5. (I) Relative protein levels of RH2B and RH2C in HCT116 cells after treatment as in C. N = 4-7. (J) Relative protein levels of RH2B and RH2C in HCT116 cells after treatment as in D. N = 5. (K) RT-qPCR analysis of *RH2A, RH2B*, *RH2C* expression in HCT116 cells after 2 h treatment with 10 μM CPT. N = 4. (L) RNase H2 activity (DRD:DNA substrate) in HT55 cells after 2 h treatment with 200 μM HU or CPT. N = 5 (1 μM CPT and 200 μM HU), 3 (10 μM CPT). (M) RNase H2 activity (DRD:DNA substrate) in HCT116 cells after 2 h treatment with CPT. N = 4. Bar graphs: Means and SEM (bars) of independent experiments are shown. Asterisks indicate p-values (ANOVA, ns: not significant, * p < 0.05, ** p < 0.01).

### Overexpression of RNASEH2B increases ribonucleotide and R-loop resolving activity

To develop tools for studying the role of increased RNase H2 activity in the response to replication stress, we generated HCT116 and BJ-hTERT cell lines for inducible overexpression of MYC-tagged RNASEH2B (Fig. 3A, B). When RNASEH2B was overexpressed in these cell lines, RNASEH2A and RNASEH2C protein levels also increased (Fig. 3B, Fig. S3A) without a corresponding increase in *RNASEH2A or RNASEH2C* mRNA expression (Fig. 3C, Fig. S3B). Co-immunoprecipitation showed that ectopic Myc-RNASEH2B interacts with endogenous RNASEH2A and RNASEH2C forming the RNase H2 complex (Fig. 3D). Ectopic RNASEH2B overexpression further increased RNase H2-specific and RNA:DNA hybrid resolution activity (Fig. 3E; Fig. S3C-F).

**Figure 3.**
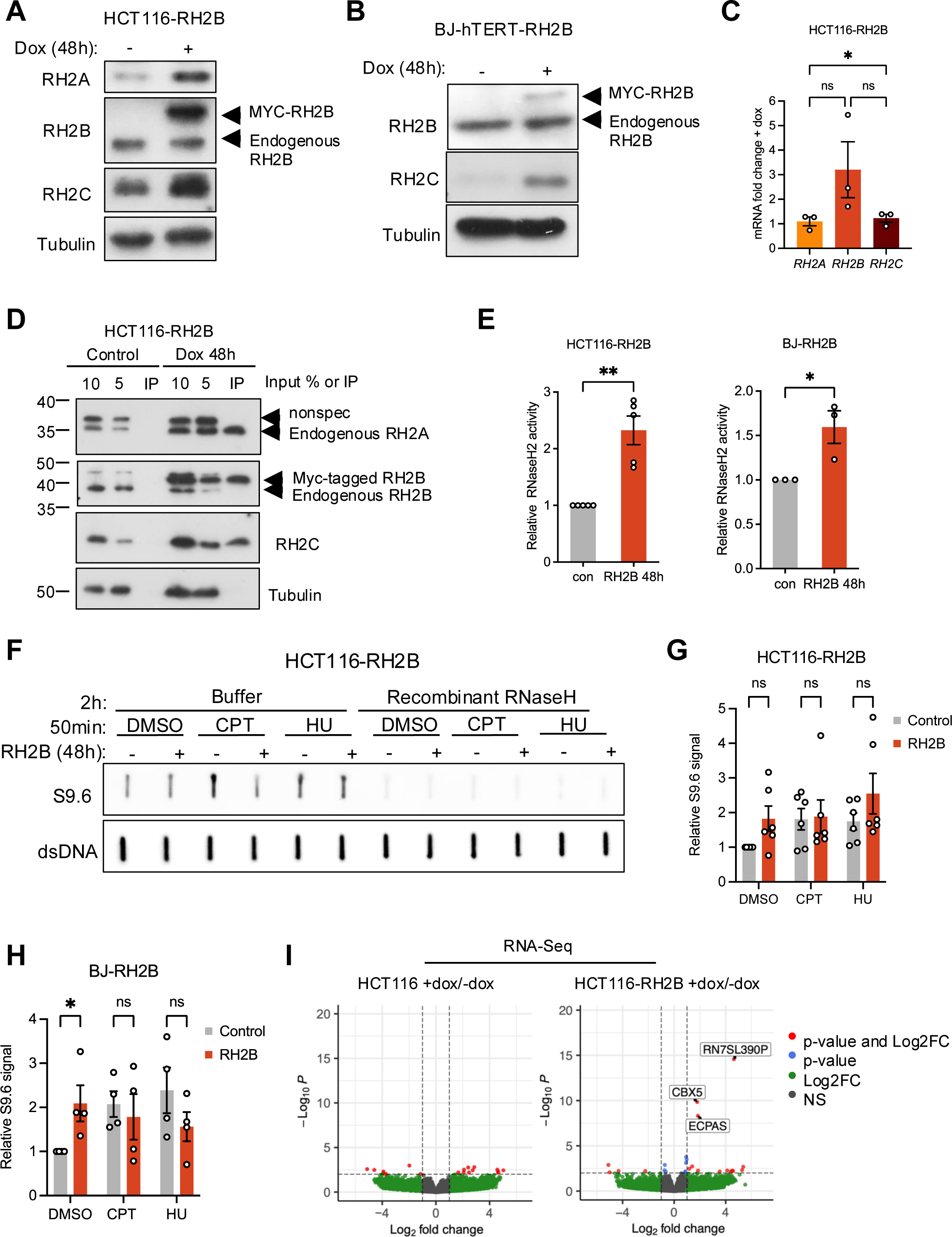
Inducible overexpression of RNASEH2B increases the levels of active RNase H2 complex. (A) Protein levels of RNASEH2B (RH2B), RNASEH2C (RH2C) and Tubulin (loading control) in BJ-hTERT-RH2B cells after +/- 48 h treatment with doxycyline (Dox). (B) Protein levels of RNASEH2A (RH2A), RH2B, RH2C and Tubulin (loading control) in HCT116-RH2B cells +/- 48 h Dox. (C) RT-qPCR analysis of *RNASEH2A (RH2A), RNASEH2B (RH2B)*, and *RNASEH2C (RH2C)* expression in HCT116-RH2B cells after 48 h Dox normalised to no Dox control. N=3. (D) Co-immunoprecipitation using MYC-trap to precipitate ectopic RH2B 48 h after RH2B induction, followed by immunoblotting for RH2A, RH2B, RH2C and Tubulin (loading control). Non-spec: non-specific band. (E) RNase H2 activity (DRD:DNA substrate and RNA:DNA substrate) in HCT116-RH2B and BJ-hTET-RH2B cells +/-48 h Dox. N=5 (HCT116), N=3 (BJ-hTERT). (F) Slot blot analysis of RNA:DNA hybrid levels in genomic DNA after 50 min treatment with 200 μM hydroxyurea (HU), 10 μM CPT or DMSO. S9.6: RNA:DNA hybrids; dsDNA: double-stranded DNA (loading control). (G) Quantification of slot blot analysis in HCT116-RH2B cells. S9.6 intensities were normalised to DMSO for each condition (control or RH2B induction). N=6. (H) Quantification of slot blot analysis in BJ-hTERT-RH2B cells as in G. N=4. (I) Volcano plot of differentially expressed genes in parental HCT116 cells treated with doxycycline (control for doxycycline effects) and HCT116-RH2B cells treated with doxycycline (RH2B overexpression). Red dots indicate differentially expressed genes with a P-value<0.01 and LogFC>2. Data from N=3. The means and SEM (bars) of independent experiments are shown. Asterisks indicate p-values (ANOVA, ns: not significant, * p < 0.05, ** p < 0.01).

To test whether increased RNase H2 activity would impact on global RNA:DNA hybrid levels, RNASEH2B overexpression was combined with a 50 min CPT or HU treatment to induce RNA:DNA hybrids. Slot blot analysis of isolated genomic DNA using the S9.6 antibody that detects RNA/DNA hybrids^46^ revealed that RNASEH2B overexpression alone increased RNA:DNA hybrid signal to some extent in untreated cells (Fig. 3F-H). CPT or HU treatment both induced around 2-fold increase in global RNA:DNA hybrid signal, but there was less or no additional increase in RNA:DNA hybrids when the drug treatments were combined with RNASEH2B overexpression (Fig. 3G, H, Fig. S4A, B). These effects were significant in BJ-hTERT fibroblasts (Fig 3H, Fig. S4B), but not in HCT116 cells (Fig 3G, Fig. S4A) where the RNA:DNA hybrid signal more variable.

RNase H2 has been shown to promote efficient transcription by interacting with RNA polymerase II^47^. Therefore, to investigate whether RNASEH2B overexpression alters the expression of cell cycle or DNA damage response genes with impact on replication stress phenotypes, we performed steady-state RNA sequencing after 48 h RNASEH2B induction in HCT116-RH2B cells. 48 h doxycycline treatment of parental HCT116 cells was used as a control (Fig. 3I). RNASEH2B overexpression changed the expression of few genes (13 genes up and 4 genes down, Fig. 3J, Table S1). Three transcripts, *EPCAS, CBX5,* and *RN7SL390P* (the latter an snRNA pseudogene embedded within *CBX5*) were significantly increased after RNASEH2B induction, but this was not reproducible by RT-qPCR (Fig. S4C, D). Gene set enrichment analysis suggested increased expression of genes involved in inflammatory responses and epithelial-mesenchymal transition, which are also responses to DNA damage or stress^48–53^, and a down-regulation of proliferation pathways (Fig. S4E, F). Taken together, while overexpression of RNASEH2B in HCT116 cells for 48 h has very little effect on expression of individual genes, subtle changes in gene expression patterns are consistent with some cellular stress in line with the increased RNA:DNA hybrid levels.

### RNASEH2B overexpression counteracts chemotherapy-induced replication fork stalling

To characterise the impact of RNASEH2B overexpression on DNA replication and replication stress, we used DNA fibre approaches. RNASEH2B overexpressing BJ-hTERT fibroblasts displayed no overt growth defects for up to 7-10 days after RNASEH2B induction (Fig. S5A-C). DNA fibre analysis using thymidine analogues chlorodeoxyuridine (CldU) and iododeoxyuridine (IdU) for 20 min each (Fig. 4A), showed that 48 h RNASEH2B overexpression caused no significant change in average fork speed (Fig. 4B) or CldU/IdU length ratios, with ratios above 1 indicating increased replication fork stalling^54^ (Fig. 4C).

**Figure 4.**
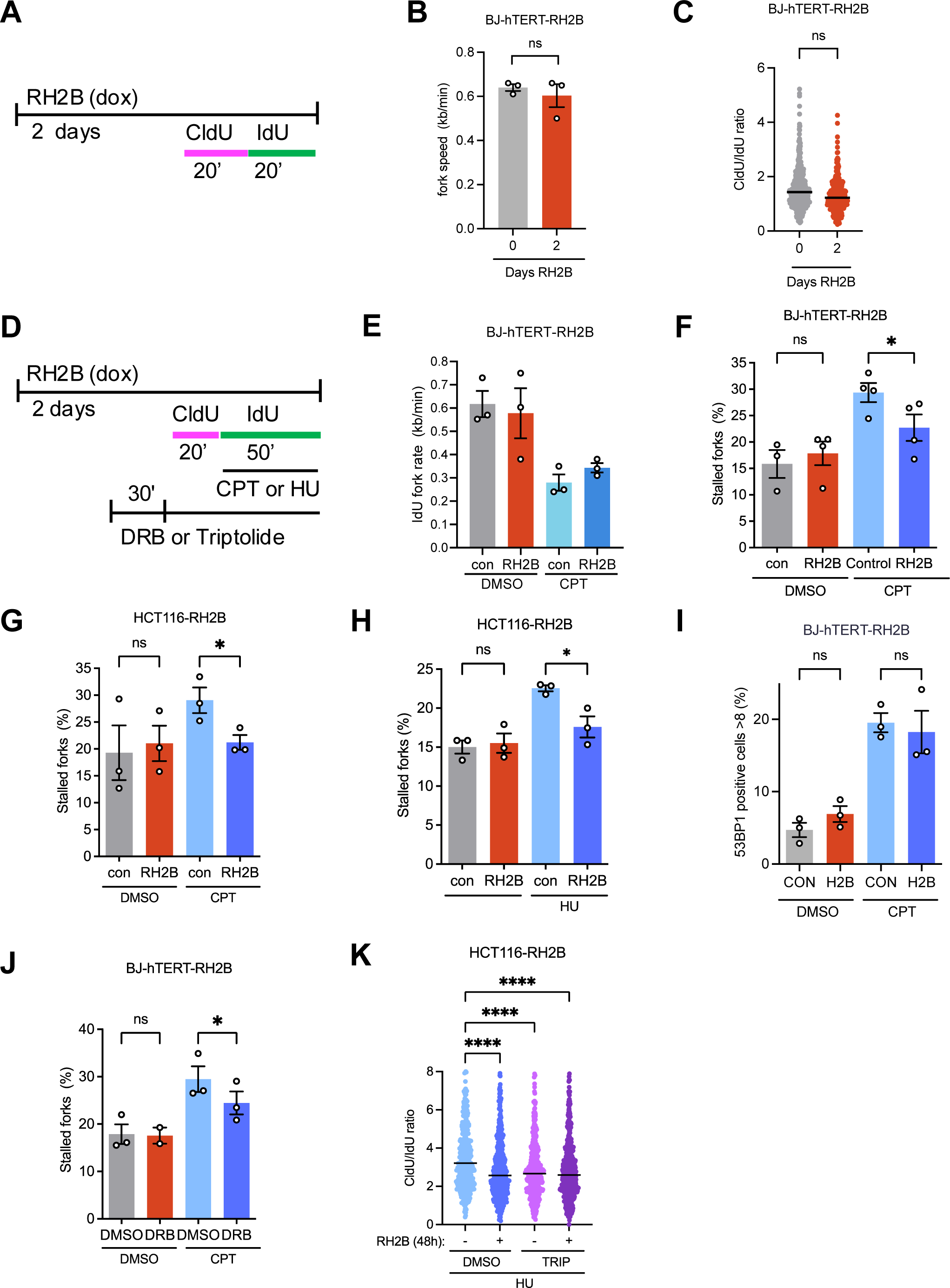
Increased RNASEH2B levels prevent drug-induced replication fork stalling. (A) DNA fibre labelling protocols after 48h RH2B overexpression. (B) Replication fork speeds in BJ-hTERT-RH2B cells +/- 48 h RH2B induction. N=3. (C) CldU/IdU ratios in BJ-hTERT-RH2B cells +/- 48 h RH2B induction. Data from 6 repeats. (D) DNA fibre labelling protocols after 48h RH2B overexpression with drug treatments. (E) Replication fork speeds (IdU label) in BJ-hTERT-RH2B cells +/- 48 h RH2B induction +/- 10 μM CPT. N=3 (F) Percentages of stalled forks (CldU labelled only) in BJ-hTERT-RH2B cells +/- 48 h RH2B induction +/- 10 μM CPT. N=4. (G) Percentages of stalled forks in HCT116-RH2B cells +/- 48 h RH2B induction +/- 100 nM CPT. CldU and IdU treatment were 30 min each. N=3. (H) Percentages of stalled forks in HCT116-RH2B cells +/- 48 h RH2B induction +/- 200 μM HU. N=3. (I) Percentages of BJ-hTERT-RH2B cells with >8 53BP1 foci +/- 48 h RH2B induction +/- 10 μM CPT. N=3. (J) Percentages of stalled forks in BJ-hTERT-RH2B cells without RH2B induction and treated with 10 μM CPT +/- 100 μM DRB. N=3. (K) CldU/IdU ratios in HCT116-RH2B cells treated with 200 μM HU +/- 48 h RH2B induction and +/- 1 μM Triptolide (TRIP). Data from 2 repeats. The means and SEM (bars) of independent experiments are shown. Asterisks indicate p-values (ANOVA or student’s t-test, ns: not significant, * p < 0.05, ** p < 0.01, **** p < 0.0001).

We then investigated the impact of RNASEH2B overexpression on chemotherapy-induced replication fork slowing and stalling, by adding CPT with the second label (IdU) for 50 min (Fig. 4D). CPT reduced IdU fork speeds by half, indicative of replication fork slowing. CPT-induced replication fork slowing was not significantly affected by RNASEH2B overexpression (Fig. 4E).

However, when quantifying stalled replication forks (fibres containing only the first or CldU label), RNASEH2B overexpression significantly reduced CPT-induced fork stalling (Fig. 4F). Similar results were obtained when using HCT116 cells instead of BJ-hTERT fibroblasts (Fig. 4G) and when treating HCT116 cells with HU instead of CPT (Fig. 4H). Therefore, RNase H2 activity counteracts CPT and HU-induced fork stalling. While fork stalling can promote fork collapse into DSBs, RNASEH2B overexpression did not affect CPT-induced DSB levels as measured by 53BP1 foci formation, with a slight but non-significant increase in 53BP1 foci after RNASEH2B overexpression alone (Fig. 4I).

It was recently reported that RNase H2 depletion decreases the degradation of HU-stalled forks and that this could be rescued with the transcription inhibitor triptolide, but not with another transcription inhibitor, DRB(Heuzé et al., 2023a). In our hands both DRB and triptolide treatments rescued CPT- or HU-induced fork stalling to a significant extent (Fig. 4J, K). This supports that in addition to promoting stalled fork resection, RNase H2 activity reduces transcription-dependent fork stalling regardless of whether the stalling is induced by CPT or HU.

### Impact of RNASEH2B overexpression on cell survival and genome stability

To investigate the impact of RNASEH2B overexpression on cell survival, we used colony formation assays. Surprisingly, RNASEH2B overexpression did not substantially affect the survival of HCT116 cells in response to CPT, HU or gemcitabine. This held true whether the treatment was administered for 24 h followed by washout (Fig. 5A-C) or continuously (Fig S6A, B). We also investigated the effect of siRNA depletion of RNASEH2A, RNASEH2B or RNASEH2C on survival. RNASEH2B depletion reduced RNASEH2-specific activity to less than 20% of control, and to the same extent as co-depletion of all subunits (Fig. 5D, E). However, depletion of any of the RNase H2 subunits did not affect the survival of HCT116 cells in response to CPT or GEM (Fig. 5F, G), with only a small reduction in survival at lower HU concentrations when RNASEH2A was depleted (Fig. 5H).

**Figure 5.**
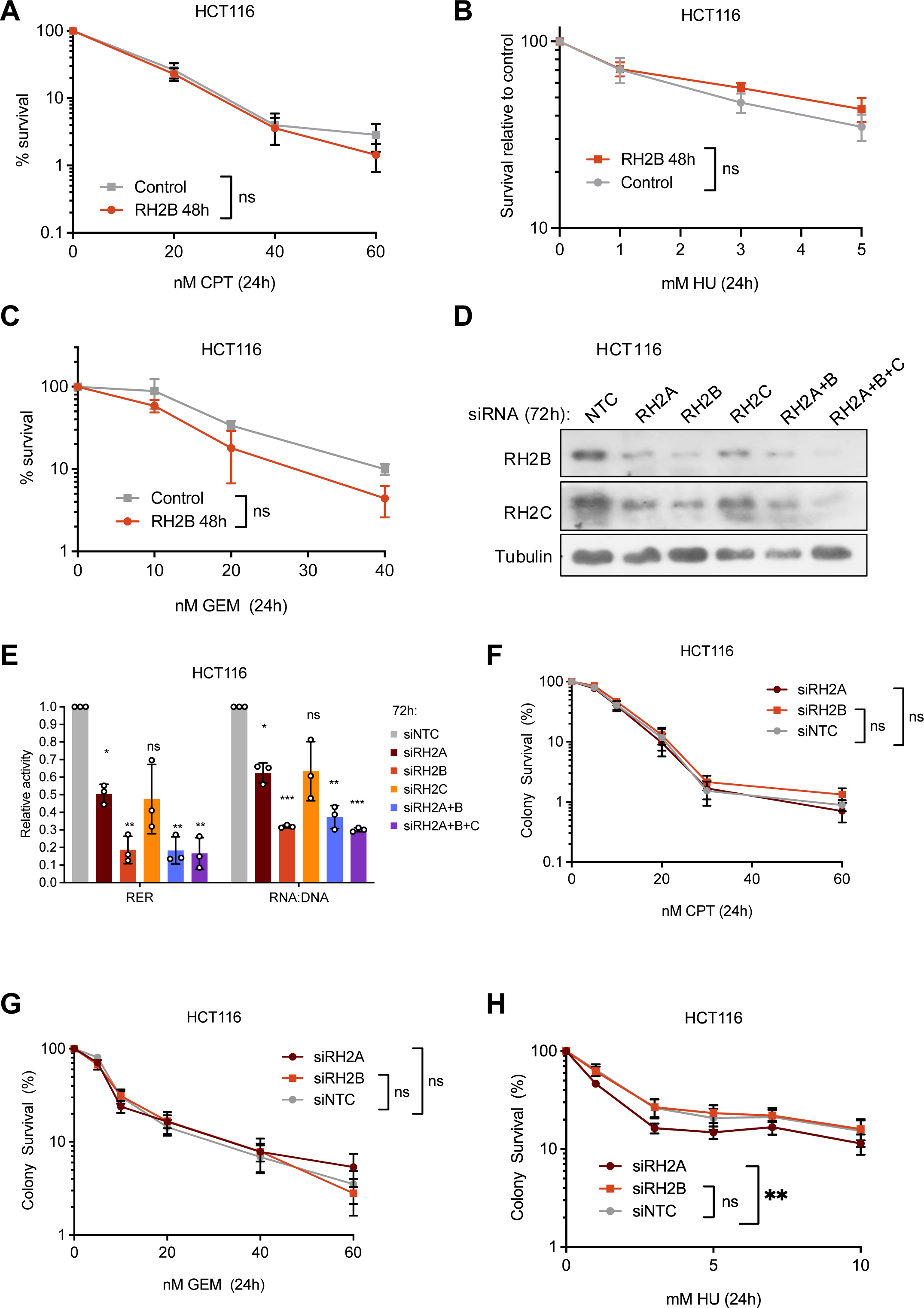
Increasing RNase H2 activity does not promote survival in response to chemotherapy. (A) Colony survival of HCT116-RH2B cells after release from 24 h CPT. N=3. (B) Colony survival of HCT116-RH2B cells after release from 24 h HU. N=6 (5 mM: N=5). (C) Colony survival of HCT116-RH2B cells after release from 24 h Gem. N =2. (D) Protein levels of RH2B, RH2C, and tubulin (loading control) in HCT116 cells after 72 h transfection with non-targeting control siRNA (siNTC) or siRNA against RH2A (siRH2A), RH2B (siRH2B) and/or RH2C (siRH2C). Cell extracts were prepared using whole cell extract buffer for RNase H2 activity assay. (E) Relative RNase H2 and RNA:DNA hybrid resolving activity in HCT116 cell extracts after 72 h transfection with non-targeting control siRNA (siNTC) or siRNA against RH2A (siRH2A), RH2B (siRH2B) and/or RH2C (siRH2C). N=3. (F) Colony survival of HCT116 cells after 72 h transfection with siNTC, siRH2B or siRH2C followed by 24 h treatment with CPT and release into fresh medium. N = 6. (G) Colony survival of HCT116 cells after 72 h transfection with siNTC, siRH2B or siRH2C followed by 24 h treatment with GEM and release into fresh medium. N = 5 (5 nM, 10 nM), N = 6 (DMSO, 20 nM, 40 nM, 60 nM). (H) Colony survival of HCT116 cells after 72 h transfection with siNTC, siRH2B or siRH2C followed by 24 h treatment with HU and release into fresh medium. N = 5. The means and SEM (bars) of independent experiments are shown. Asterisks indicate p-values (ANOVA, ns: not significant, * p < 0.05, ** p < 0.01, *** p < 0.001).

Rather than promoting cell survival, RNASEH2B overexpression may protect against sub-lethal genome instability and/or prevent innate immune activation. To investigate the impact of RNASEH2B overexpression on genome instability, we quantified micronucleus formation as a marker of genome instability in response to short-term treatment with CPT, GEM or HU in HCT116 cells (Fig. 6A). RNASEH2B overexpression prevented HU-induced micronucleus formation (Fig. 6B; Fig. S6C), had a small effect on reducing micronuclei in response to GEM (Fig. 6C; Fig. S6D), but no effect on the induction of micronuclei in response to CPT (Fig. 6D; Fig. S6E). In BJ-hTERT cells, we observed micronucleus induction only after 24 h continuous drug treatment and under these conditions the effect of RNASEH2B overexpression was less pronounced (Fig. S6F-H).

**Figure 6.**
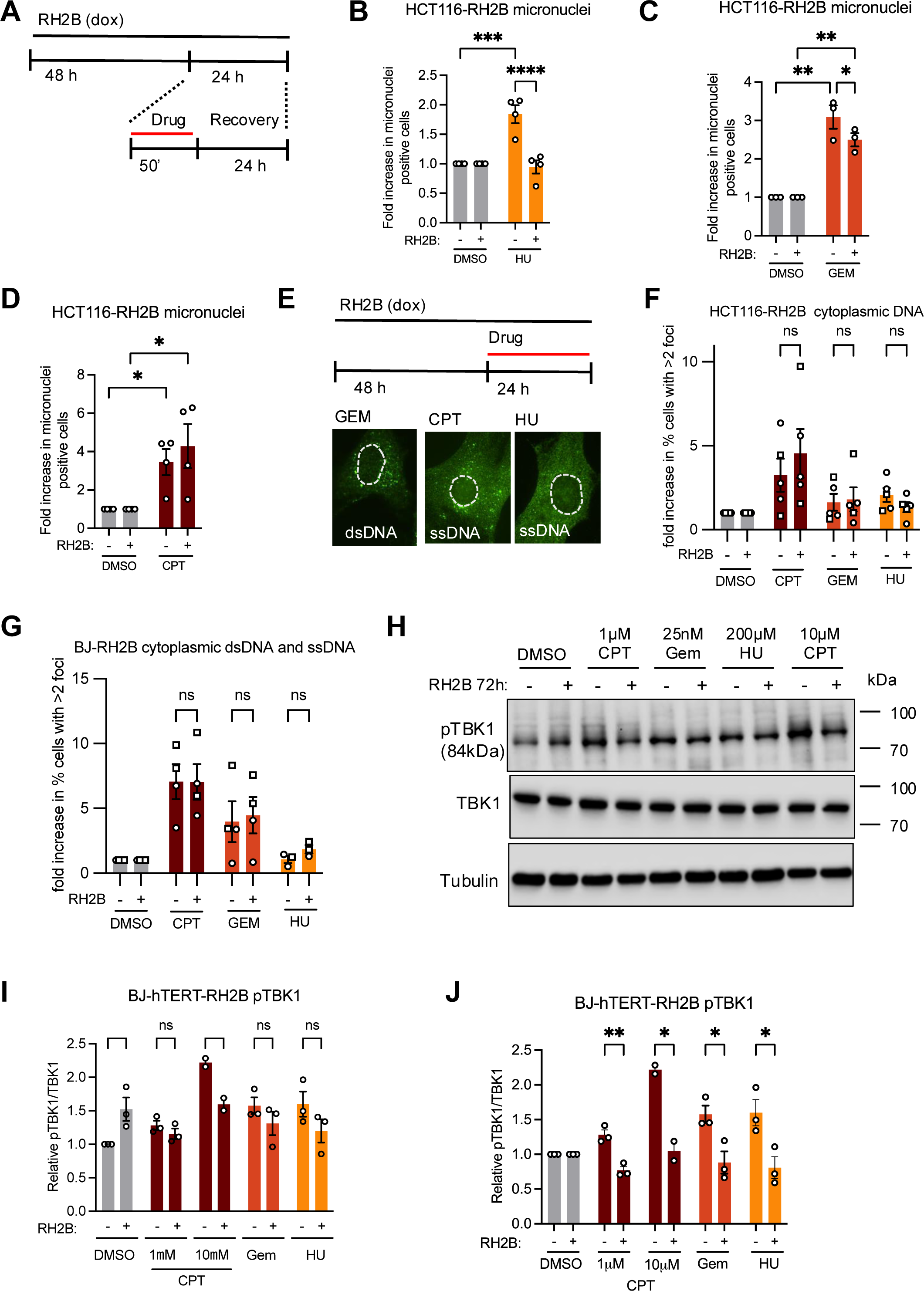
Increasing RNase H2 activity can reduce genome instability and cytoplasmic nucleic acid signalling. (A) Treatment schematic for micronucleus quantification in HCT116 cells. (B) Fold increase in HCT116-RH2B cells with micronuclei +/- RH2B induction +/- 2mM HU. N=4. (C) Fold increase in HCT116-RH2B cells with micronuclei +/- RH2B induction +/- 1 μM GEM. N=3. (D) Fold increase in HCT116-RH2B cells with micronuclei +/- RH2B induction +/- 10 μM CPT. N=3. (E) Treatment schematic and representative images for cytosolic DNA staining after drug treatments. (F) Fold increase in percentages of HCT116-RH2B cells with cytoplasmic ssDNA or dsDNA staining +/- 48 h RH2B induction and 24 h treatment with 1 μM CPT, 25 nM GEM, 200 μM HU or DMSO. Circles: dsDNA antibody, squares: ssDNA antibody. N=5. (G) Fold increase in percentages of BJ-hTERT-RH2B cells with cytoplasmic ssDNA or dsDNA staining +/- 48 h RH2B induction and 24 h treatment with 1 μM CPT, 25 nM GEM, 200 μM HU or DMSO. Circles: dsDNA antibody, squares: ssDNA antibody. N=4 (CPT and GEM), N=3 (HU). (H) Protein levels of phospho-S172 TBK1, TBK1 and Tubulin (loading control) in BJ-RH2B cells +/- 48 or 72 h RH2B induction and 48 h treatment with drugs or DMSO as indicated. (I) Relative protein levels of phospho-TBK1 normalised to TBK1 in BJ-RH2B cells after treatment as in H. N=3, N=2 (10 μM CPT). (J) Relative protein levels of phospho-TBK1 as in I, normalised to DMSO also for samples with RH2B induction. The means and SEM (bars) of independent experiments are shown. Asterisks indicate p-values (ANOVA, ns: not significant, * p < 0.05, ** p < 0.01, *** p < 0.001, **** p < 0.0001).

Replication stress can lead to accumulation of nucleic acids in the cytoplasm, including RNA:DNA hybrids, that can potentially activate innate immune signalling pathways^12^. DNA damage could similarly activate this pathway through accumulation of micronuclear DNA^48–50^ or other sources of dsDNA^52,55^ or ssDNA^51^. We also investigated other forms of cytosolic DNA using antibodies against single- or double-stranded DNA. We found that 24 h treatment with CPT or GEM resulted in foci of cytoplasmic DNA as measured by antibodies raised against either single- or double-stranded DNA (Fig. 6E), but that the levels of these foci were unaffected by RNASEH2B overexpression in HCT116 and BJ-hTERT cells (Fig. 6F, G; Fig. S6I, J). Because HCT116 cells are cGAS-STING signalling-deficient^56^, we used BJ-hTERT-RH2B cells to investigate the impact of RNASEH2B overexpression on cytoplasmic nucleic acid signalling pathways. We quantified autophosphorylation of tank binding kinase 1 (TBK1), a serine/threonine kinase directly activated by several nucleic acid sensing pathways including cGAS-STING and RIG-I, which then regulates innate immunity responses including activation of type I interferon and interferon-stimulated genes (ISGs)^57^. Activating TBK1 autophosphorylation on Serine 172 was increased both after 72 h RNASEH2B overexpression and by 48 h drug treatments in uninduced BJ-hTERT-RH2B cells and there was no further increase when drugs were combined with RNASEH2B overexpression (Fig. 6H-J). 72 h RNASEH2B overexpression does not increase micronuclei formation or cytoplasmic DNA foci, but increases RNA:DNA hybrid signal, at least in BJ-hTERT cells (Fig. 3H, Fig. S4B). Our results are therefore consistent with RNASEH2B overexpression affecting the levels of TBK1 activation, but this may be through an impact on RNA-containing nucleic acid species such as RNA:DNA hybrids^12^.

### RNASEH2B counteracts HRAS-induced replication fork stalling and cell death

To investigate the role of RNase H2 upregulation in response to RAS activation, we combined HRAS^G12V^ induction with siRNA depletion of RNASEH2B to counteract the RAS-induced increase (Fig. 7A, B). As reported previously, HRAS^G12V^ strongly decreased replication fork speeds^6^, however, RNASEH2B depletion did not decrease fork speeds further (Fig. 7C). Instead, RNASEH2B depletion increased CldU/IdU length ratios specifically in response to HRAS^G12V^ induction, suggesting increased replication fork stalling when preventing HRAS^G12V^-driven RNase H2 upregulation (Fig. 7D). Combining RNASEH2B depletion with HRAS^G12V^ induction led to no additional increase in γH2AX intensity or 53BP1 foci formation compared to HRAS^G12V^ induction alone (Fig. 7E, F). Oncogenic HRAS is known to initially increase proliferation, but then cause growth arrest due to apoptosis and senescence^58^. After HRAS^G12V^ induction, cell numbers increase for 3 days, while growth stops between 6 and 8 days^4–6^. To investigate the impact of RNase H2 on cell growth in presence of HRAS^G12V^, we measured proliferation by cell counting over a 7-day period after HRAS^G12V^ induction with or without depleting RNASEH2B using siRNA. RNASEH2B siRNA transfection was maintained for 8 days for all samples, while the length of HRAS^G12V^ induction was varied (Fig. 7G). While there was no difference in HRAS^G12V^-induced proliferation up to day 3 when HRAS^G12V^ was combined with RNASEH2B depletion, at day 7 this led to an additional decrease in cell numbers (Fig. 7H, Fig. S7A). Flow cytometry analysis revealed increased percentages of cells with sub-G1 DNA content when HRAS^G12V^ was combined with RNASEH2B depletion, indicating increased cell death (Fig. 7I, Fig. S7B). This suggests that RNase H2 activity is required to reduce replication stress for longer-term cell proliferation and survival in the presence of oncogenic RAS.

**Figure 7.**
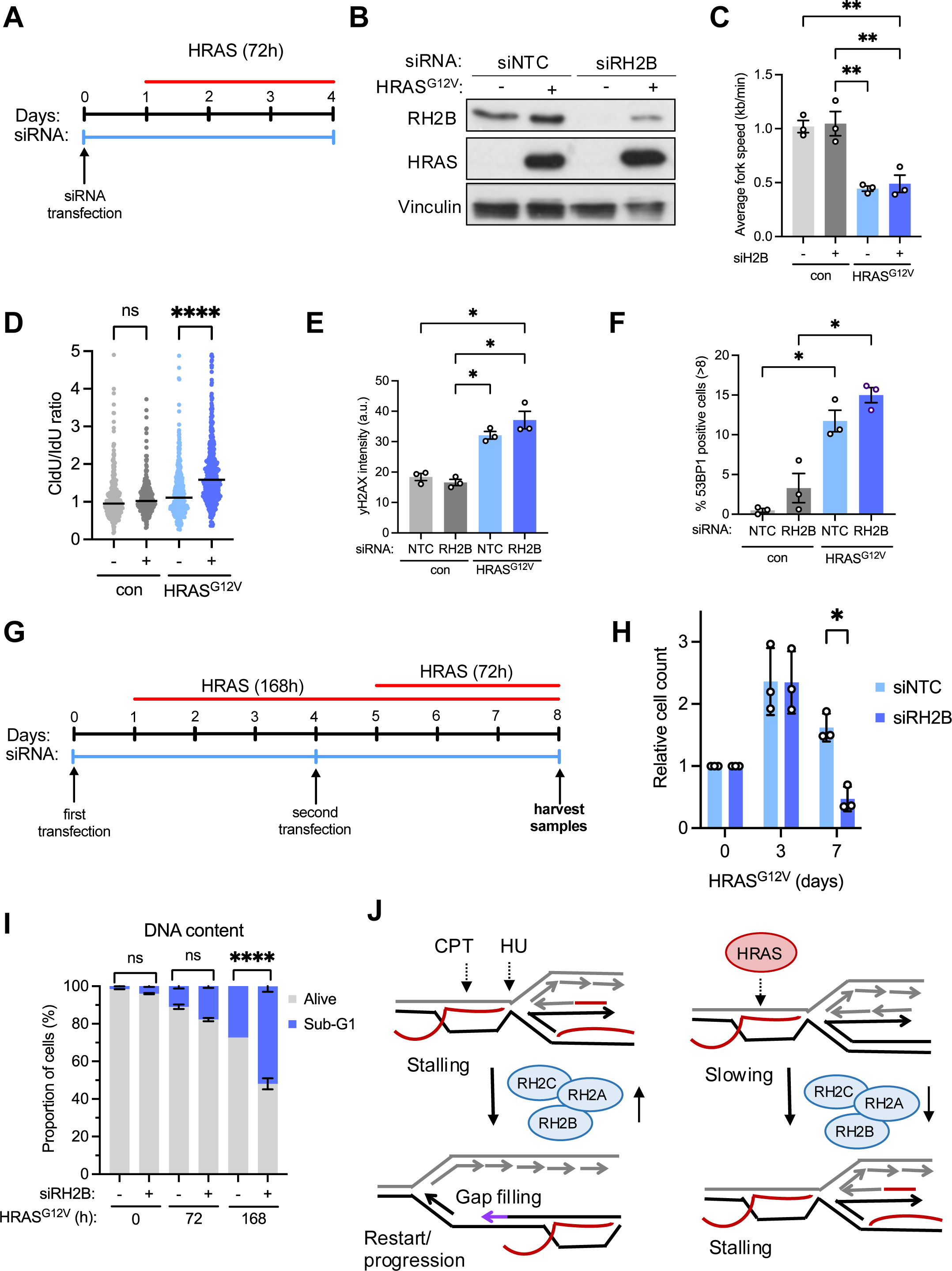
Oncogenic HRAS-induced increase in RNase H2 activity prevents replication fork stalling and cell death. (A) BJ-hTERT HRAS^V12ERTAM^ cells were transfected for 96 h with non-targeting control siRNA (siNTC) or RH2B siRNA (siRH2B) and HRAS^G12V^ was induced after 24 h for 72 h. (B) Protein levels of RNASEH2B (RH2B), HRAS^G12V^ and Vinculin (loading control) after treatment as in A. (C) Replication fork speeds +/- 72 h HRAS^G12V^ induction in presence of siNTC or siRH2B. N=3. (D) CldU/IdU ratios +/- 96 h siNTC or siRH2B and +/- 72 h HRAS^G12V^ induction. CldU and IdU treatment were 20 min each (con) and 30 min each (HRAS^G12V^). Data from 3 repeats. (E) Mean intensities of yH2AX staining +/- 72 h HRAS^G12V^ induction in presence of siNTC or siRH2B. Data from N=3. (F) Percentages of cells with >8 53BP1 foci +/- 72 h HRAS^G12V^ induction in presence of siNTC or siRH2B. N= 3. (G) Strategy for long-term HRAS^G12V^ induction with RNASEH2B depletion. (H) Relative proliferation after 8 days transfection with non-targeting control or RH2B siRNA and HRAS^G12V^ induction for the times indicated. Cell counts for siNTC or siRH2B were each normalised to no HRAS^G12V^ induction (0). N = 3. (I) Percentages of sub-G1 (dead) versus alive cells after 8 days transfection with siNTC (-) or RH2B siRNA (+) and HRAS^G12V^ induction for the times indicated, measured by flow cytometry with propidium iodide. N=2 (siNTC 186 h N=1). (J) Model: Oncogenic HRAS^G12V^, CPT and HU cause RNA:DNA hybrids (red) ahead of or behind the fork. CPT and HU-induced RNA:DNA hybrids stall forks unless RNase H2 is overexpressed. RNase H2 overexpression potentially comes at the expense of increased RNA:DNA hybrids elsewhere in the genome. HRAS^G12V^-induced RNA:DNA hybrids stall forks only when RNase H2 is depleted. Bar graphs: Means and SEM (bars) of independent experiments are shown. Scatter plots: Medians (lines) are shown. Asterisks indicate p-values (ANOVA, ns: not significant, * p < 0.05, ** p < 0.01, **** p < 0.0001).

## Discussion

We have shown that RNase H2 protein levels increase in response to replication stress from different sources including oncogenes and cancer chemotherapy drugs. RNase H2 levels may be regulated transcriptionally, but possibly also post-transcriptionally in response to replication stress. RNASEH2B overexpression increases RNase H2 activity and prevents chemotherapy- and oncogene-induced replication fork stalling, with the potential to impact on innate immune sensing of nucleic acids or cell survival.

### Co-regulation of RNASEH2B and other RNase H2 subunits

To model increased RNase H2 protein levels, we chose to focus on RNASEH2B. Although not the catalytic subunit, RNASEH2B is required for the nuclear localisation of the RNase H2 complex^17^ and interacts with the DNA sliding clamp, PCNA, which could localise it to replication forks and sites of DNA repair^16^. We occasionally observed that RNASEH2B was up-regulated while RNASEH2C remained constant, such as in response to Cyclin E overexpression. RNASEH2B appears to have a crucial role in regulating localisation and function of the RNase H2 complex and RNASEH2B depletion or knockout has been repeatedly used to reduce RNase H2 activity^25,26,31^, which was also the most effective strategy in our hands.

Our data agree with observations from knockout models showing that the three RNase H2 subunits are co-regulated at the protein level. *RNASEH2B* knockout or mutation can destabilise the complex via loss or reduction of both RNASEH2A and RNASEH2C, respectively^14,15^. Similarly, RNASEH2B overexpression can increase RNASEH2C, and vice versa. RNASEH2B overexpression is sufficient to increase cellular RNase H2 activity (Fig 3E). Therefore, importantly, phenotypic changes resulting from RNASEH2B overexpression currently do not imply specific non-RNase H2-related functions of this subunit.

### Impact on RNA:DNA hybrids and transcription

Surprisingly, RNASEH2B overexpression appeared to increase rather than decrease global RNA:DNA hybrid levels. It was previously reported that RNASEH2A depletion did not increase global RNA:DNA hybrid levels ^31^. The impact of altering RNase H2 activity only global hybrid levels is therefore not straightforward. Excessive RNase H2 activity may increase hybrid levels by disrupting RNA:DNA hybrid homeostasis through RNase H2 functions from regulating transcription to DSB repair^59,60^. We observed a small increase in levels of 53BP1 foci, a marker of DSB formation, in cells overexpressing RNASEH2B (Fig. 4I). RNASEH2B overexpression could also increase binding to interaction partners such as PCNA^14^, which might disrupt DNA replication and DNA repair processes that are regulated by PCNA. Other putative RNase H2 interactors are involved in transcriptional regulation^61^. While there was no overt impact of RNASEH2B overexpression on global nascent RNA synthesis or gene expression, there could be subtler changes at RNA:DNA hybrid hotspots such as promoter and terminator regions of genes^3^.

### RNASEH2B overexpression rescues drug-induced replication fork stalling

RNASEH2B overexpression rescued a subset of CPT- and HU-induced replication stress, specifically replication fork stalling, potentially via reducing TRCs or RNA:DNA hybrids at replication forks. This complements previous reports in yeast and human cells using RNase H2 loss or depletion to show that RNase H2 promotes nascent strand degradation in presence of HU or CPT, and replication fork progression in presence of HU, by removing RNA:DNA hybrids derived either from ongoing transcription or from Okazaki primers^30,31^. Our data expand on these findings by directly showing that RNase H2 counteracts replication fork stalling in human cells. Our data are consistent with the idea that RNase H2 supports replication fork restart by promoting nascent strand degradation, as acute transcription inhibition both promoted nascent strand degradation^31^ and prevented fork stalling (this study). Our data also support that in addition to nucleotide depletion, transcription is involved during fork stalling induced by HU. In line with this, HU also increases reactive oxygen species and causes fork stalling that can be alleviated by removing RNA:DNA hybrids^62^.

There are still open questions regarding the role of RNase H2 at stalled replication forks. Increased RNase H2 activity has been proposed to interfere with fork restart after HU or aphidicolin if RNase H2 localisation to replication forks is not properly controlled by factors such as the proteasome shuttle protein DDI1/2^63^. We also observed that RNASEH2B overexpression resulted in some signs of stress, especially increased RNA:DNA hybrid levels. The molecular mechanisms underlying both HU- or CPT-induced replication fork stalling and their relationship to RNA:DNA hybrids and nascent strand degradation require further investigation. The DNA fibre labelling protocol used in our study (Fig. 5D) could theoretically lead to forks resected during drug treatments appearing as stalled forks. However in that case, the promotion of resection by RNase H2 would lead to increased, not decreased, fork stalling when RNASEH2B is overexpressed.

### Effects of RNASEH2B overexpression on genome stability and cell survival

Even though RNase H2 activity is implicated not only in replication stress responses but also in DSB repair^59,60^, RNASEH2B overexpression had little effect on CPT-induced DSB levels (Fig. 4I) or colony survival in response to CPT, GEM or HU in HCT116 cells. It was previously reported that constitutive RNASEH2A overexpression protected LNCaP prostate cancer cells against CPT or etoposide^34^ and that RNase H2 depletion increased HU sensitivity in HeLa cells^15^. In contrast, RNASEH2A depletion was shown to decrease HU sensitivity in U2OS cells^63^. Finally, RNase H2 deficient budding yeast are more sensitive to replication stress-inducing drugs^64^. Altogether, more remains to be discovered about the precise impact of RNase H2 levels on drug sensitivity in human cells.

Even if replication stress response factors do not promote survival, they can be important to safeguard genome stability in surviving cells. RNASEH2B overexpression reduced micronuclei formation in response to HU treatment. It is possible that the extent of micronucleus induction by CPT and GEM matched the extent of DSB formation^65^ more than levels of fork stalling.

We observed phosphorylation of TBK1 in BJ-hTERT cells in response to drug treatments, indicating activation of cytoplasmic nucleic acid sensing pathways. Our data do not rule out that some of the observed TBK1 activation after drug-induced replication stress could be linked to formation of micronuclei or other cytosolic DNA species. Overall, TBK1 activation, both in response to RNASEH2B overexpression and to drug treatments, correlated best with nuclear RNA:DNA hybrid formation.

### RNase H2 prevents RAS-induced replication fork stalling

As with the response to replication stress-inducing drugs, RNase H2 specifically alleviates fork stalling rather than fork slowing induced by HRAS^G12V^. This suggests that HRAS^G12V^-induced transcription and/or RNA:DNA hybrids can cause fork stalling through a mechanism that is similar to the fork stalling induced by HU or CPT. In contrast to HU or CPT, HRAS^G12V^ only causes fork stalling when RNase H2 is also depleted. This suggests that the increase in RNase H2 levels after HRAS^G12V^ induction has a similar rescuing effect as ectopic RNase H2 overexpression, or fork stalling may require accumulation of higher levels of RNA:DNA hybrids, if there is no additional nucleotide depletion^6^ or DNA damage (Figure 7J). Again, the relationship of fork stalling to nascent strand degradation requires further investigation. RNase H2 depletion accelerated growth arrest and cell death in response to oncogenic HRAS. This complements previous reports that down-regulation of other DNA repair and - replication proteins, such as BRIP1 (involved in BRCA1-mediated DSB repair) and RRM2 (a ribonucleotide reductase subunit) accelerates HRAS-induced senescence or growth arrest^66,67^, presumably by further exacerbating HRAS-induced replication stress. Our data suggest that oncogenic HRAS-expressing cells rely on RNase H2 to keep replication stress at tolerable levels for proliferation and survival.

In conclusion, increased levels of RNase H2 subunits may be indicators of the response to chemotherapy or oncogene-induced replication stress, warranting further investigation. RNase H2 mRNA levels are increased in some CIN+ colon cancers, and it is possible that protein levels are also be up-regulated post-transcriptionally. Deciphering the mechanisms that regulate RNase H2 protein levels, activity and localisation in response to cellular stress will contribute to our understanding of replication stress responses in cancer.

## Acknowledgements

R.J.W., C.W. and R.D.W.K. were supported by a Cancer Research UK Programme Foundation award to E.P. and Ad.K. (C25526/A28275). P.K. was supported by a Worldwide Cancer Research grant (13-1048) to E.P. We thank Dr Angelo Agathanggelou for technical help.

## Author contributions

E.P., C.H.M.T. and P.K. conceived the study; R.J.W., Ab.K., S.A.P. and R.D.W.K. performed experiments; R.J.W., Ab.K., P.K., M.A.M.R. and E.P. designed experiments; C.W. and Ad.K. performed and supervised RNA-seq data analysis; R.J.W. and E.P. wrote the paper.

## Competing Interests

The authors declare that they have no competing financial interest.

## Data Availability Statement

The authors declare that all the data supporting the findings of this study are available within the article and its supplementary information files and from the corresponding authors upon reasonable request. The datasets generated during and analysed during the current study are available in the GEO repository, https://www.ncbi.nlm.nih.gov/geo/query/acc.cgi?acc=GSE276623.

## Notes

https://www.ncbi.nlm.nih.gov/geo/query/acc.cgi?acc=GSE276623

### Competing Interest Statement

The authors have declared no competing interest.

