## Supplemental Data for "Human RNase H2 upregulation counteracts oncogene- and chemotherapy-induced replication stress"

### Supplementary Figures

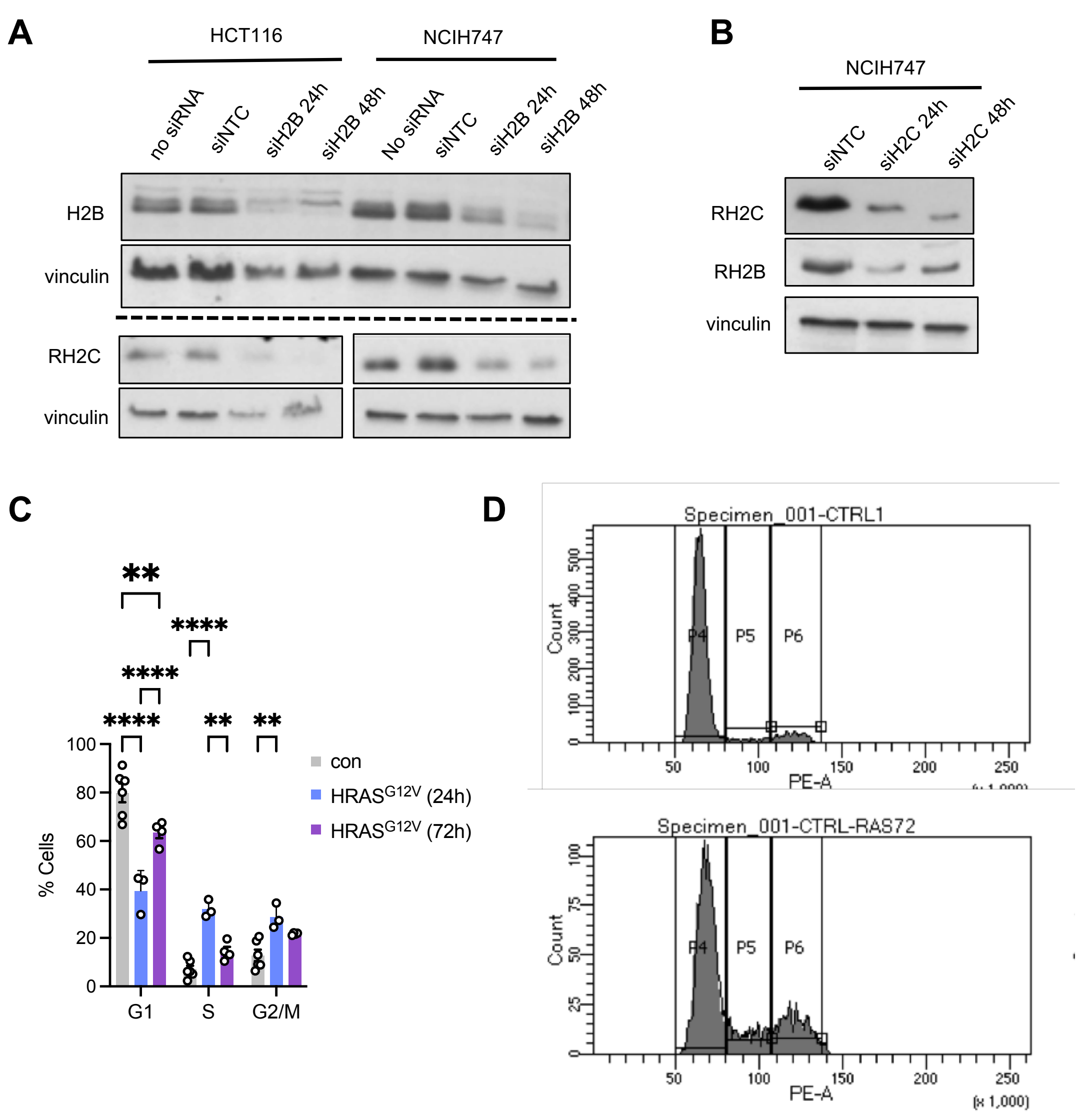


**Figure S1. Validation of RNaseH2 subunit antibodies**
**(A)** Protein levels of RNaseH2B (RH2B), RNaseH2C (RH2C), and vinculin (loading control) in HCT116 or NCIH747 cells after 24 or 48 h transfection with non-targeting control siRNA (siNTC) or RH2B siRNA (siRH2B), or no transfection (no siRNA). **(B)** Protein levels of RH2B and RH2C and vinculin (loading control) in NCIH747 cells after 24 or 48 h transfection with non-targeting control siRNA (siNTC) or RH2C siRNA (siRH2C).
**(C)** Quantification of cell cycle distribution in BJ-HRAS$^{G12V}$ cells with and without 24 or 72 h HRAS$^{G12V}$ induction, based on measuring DNA content using propidium iodide staining and flow cytometry. N=3 (24h), N=4 (72h), N=6 (control). **(D)** Flow cytometry gating strategy for quantification of cell cycle distribution as in D. Cells were stained with propidium iodide.
The means and SEM (bars) of independent experiments are shown. Asterisks indicate p-values (ANOVA, * $p < 0.05$, * $p < 0.05$, ** $p < 0.01$, **** $p < 0.0001$).

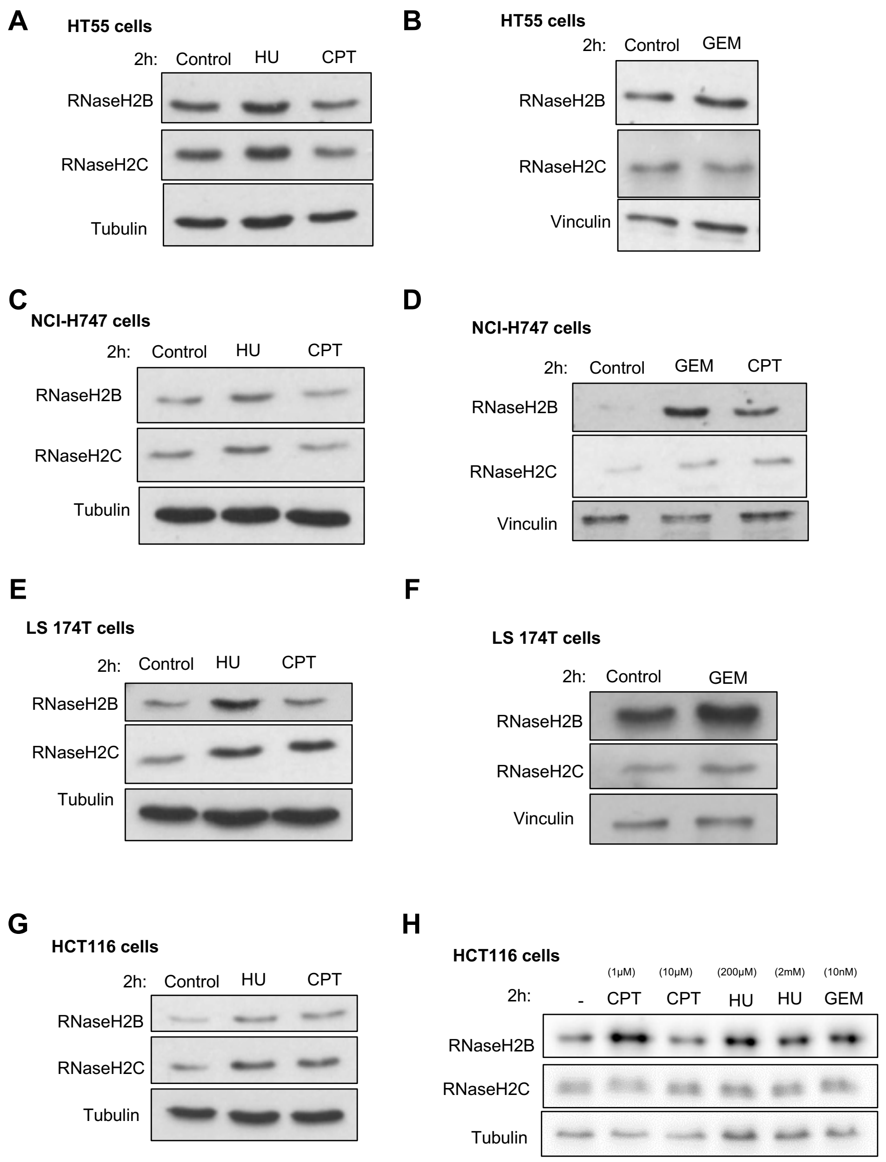


**Figure S2. Representative Western blots of RNaseH2 subunit levels after chemotherapy treatment in colon cancer cell lines**
**(A)** Protein levels of RH2B, RH2C and tubulin (loading control) in HT55 cells after 2 h treatment with 200 μM hydroxyurea (HU) or 10 μM camptothecin (CPT). **(B)** Protein levels of RH2B, RH2C and vinculin (loading control) in HT55 cells after 2 h treatment with 25 nM gemcitabine (GEM). **(C)** Protein levels of RH2B, RH2C and vinculin (loading control) in NCIH747 cells after 2 h drug treatment with 200 μM HU or 10 μM CPT. **(D)** Protein levels of RH2B, RH2C and vinculin (loading control) in NCIH747 cells after 2 h treatment with 25 nM GEM or 10 μM CPT. **(E)** Protein levels of RH2B, RH2C and vinculin (loading control) in LS174T cells after 2 h treatment with 200 μM HU or 10 μM CPT. **(F)** Protein levels of RH2B, RH2C and vinculin (loading control) in LS174T cells after 2 h treatment with 25 nM GEM. **(G)** Protein levels of RH2B, RH2C and tubulin (loading control) in HCT116 cells after 2 h treatment with 200 μM HU or 10 μM CPT. **(H)** Protein levels of RH2B, RH2C and tubulin (loading control) in HCT116 cells after 2 h treatment with HU, CPT or GEM as indicated.

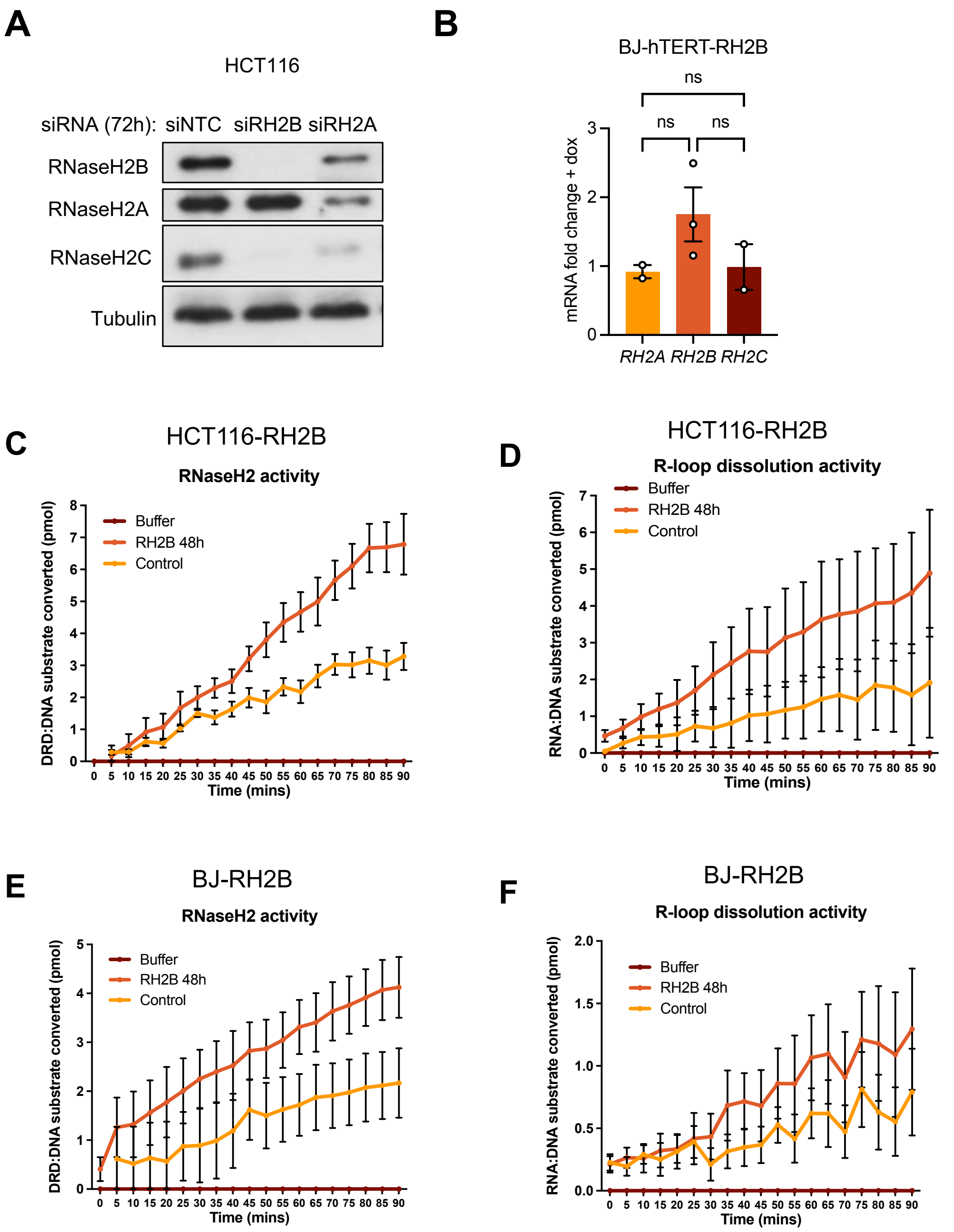


**Figure S3. Validation of RNaseH2A antibody, and RNaseH2 activity assays.** **(A)** Protein levels of RNaseH2B (RH2B), RNaseH2A (RH2A), RNaseH2C, and Tubulin (loading control) in HCT116 cells 72 h after transfection with non-targeting control siRNA (siNTC), siRNA against RH2B (siRH2B) or R2HA (siRH2A). **(B)** RT-qPCR analysis of *RNASEH2A (RH2A), RNASEH2B (RH2B)*, and *RNASEH2C (RH2C)* expression in BJ-hTERT-RH2B cells after 48 h RH2B induction compared to no dox control. N=3. **(C)** DRD:DNA substrate converted over time during incubation with buffer only, whole cell extract from HCT116-RH2B cells with (RH2B 48h) or without (control) 48 h RH2B induction. N=4 **(D)** DRD:DNA substrate converted over time during incubation with buffer, whole cell extract from HCT116-RH2B cells with (RH2B 48h) or without (control) 48 h RH2B induction. N=2. **(E)** RNA:DNA substrate converted over time during incubation with buffer only, whole cell extract from BJ-RH2B cells with (RH2B 48h) or without (control) 48 h RH2B induction. N=4 **(F)** RNA:DNA substrate converted over time during incubation with buffer, whole cell

extract from BJ-RH2B cells with (RH2B 48h) or without (control) 48 h RH2B induction. N=3.
The means and SEM (bars) of independent experiments are shown. Asterisks indicate p-values (ANOVA, ns: not significant).

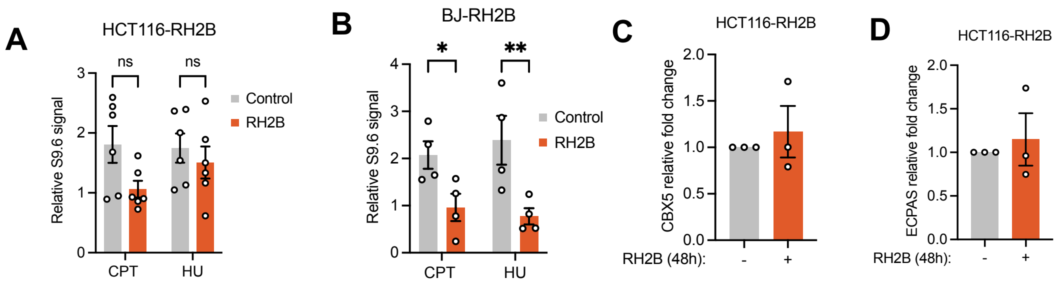


**E**

| | Gene set (UP) | # Genes in set | Enrichment score | Normalised enrichment score | NOM p-val | FDR q-val |
|---|---|---|---|---|---|---|
| 1 | Interferon alpha response | 97 | 0.62 | 1.63 | 0 | 0.009 |
| 2 | Mitotic spindle | 199 | 0.55 | 1.55 | 0 | 0.017 |
| 3 | Interferon gamma response | 200 | 0.51 | 1.47 | 0.002 | 0.045 |
| 4 | Hedgehog signalling | 36 | 0.63 | 1.41 | 0.034 | 0.064 |
| 5 | Apical surface | 44 | 0.56 | 1.31 | 0.082 | 0.171 |
| 6 | Epithelial mesenchymal transition | 200 | 0.46 | 1.29 | 0.024 | 0.183 |
| 7 | Protein secretion | 96 | 0.47 | 1.24 | 0.061 | 0.24 |
| 8 | Inflammatory response | 200 | 0.41 | 1.19 | 0.084 | 0.365 |

**F**

| | Gene set (DOWN) | # Genes in set | Enrichment score | Normalised enrichment score | NOM p-val | FDR q-val |
|---|---|---|---|---|---|---|
| 1 | Myc targets V1 | 200 | -0.74 | -2.21 | 0 | 0 |
| 2 | Oxidative phosphorylation | 200 | -0.69 | -2.07 | 0 | 0 |
| 3 | Myc targets V2 | 58 | -0.76 | -1.92 | 0 | 0.001 |
| 4 | DNA repair | 150 | -0.57 | -1.66 | 0 | 0.003 |
| 5 | E2F targets | 200 | -0.55 | -1.66 | 0 | 0.003 |
| 6 | MTORC1 signalling | 200 | -0.49 | -1.45 | 0 | 0.03 |
| 7 | Unfolded protein response | 113 | -0.52 | -1.44 | 0.003 | 0.026 |
| 8 | UV response UP | 158 | -0.48 | -1.4 | 0 | 0.036 |
| 9 | Reactive oxygen species pathway | 49 | -0.56 | -1.39 | 0.036 | 0.034 |
| 10 | p53 pathway | 200 | -0.44 | -1.31 | 0.022 | 0.07 |
| 11 | Fatty acid metabolism | 158 | -0.43 | -1.25 | 0.038 | 0.107 |
| 12 | Adipogenesis | 200 | -0.4 | -1.22 | 0.032 | 0.129 |
| 13 | TNFA signalling via NFKB | 200 | -0.4 | -1.19 | 0.059 | 0.157 |
| 14 | G2M checkpoint | 200 | -0.39 | -1.15 | 0.099 | 0.201 |

**Figure S4. Impact of RNaseH2B overexpression on RNA:DNA hybrid levels and gene expression.**
**(A)** Quantification of slot blot analysis of RNA:DNA hybrid levels in genomic DNA from BJ-RH2B cells after 50 min treatment with 200 μM hydroxyurea (HU), 10 μM CPT or DMSO as in Fig. 3F, without normalisation to RH2B DMSO. N=4. **(B)** Quantification of slot blot analysis of RNA:DNA hybrid levels in genomic DNA from HCT116-RH2B cells after 50 min treatment with 200 μM hydroxyurea (HU), 10 μM CPT or DMSO as in Fig. 3F, without normalisation to RH2B DMSO. N=6. **(C)** RT-qPCR analysis of *CBX5* expression in HCT116-RH2B cells after 48 h RNH2B induction. N=3. **(D)** RT-qPCR analysis of *EPCAS* expression in HCT116-RH2B cells after 48 h RNH2B induction. N=3. **(E)** Top enriched gene sets that are specifically upregulated upon 48 h dox treatment in inducible HCT116-RH2B cells, but not in parental controls. **(F)** Top enriched gene sets that specifically downregulated upon 48 h dox treatment in inducible HCT116-RH2B cells, but not in parental controls.
The means and SEM (bars) of independent experiments are shown. Asterisks indicate p-values (ANOVA, ns: not significant, * $p < 0.05$, ** $p < 0.01$).

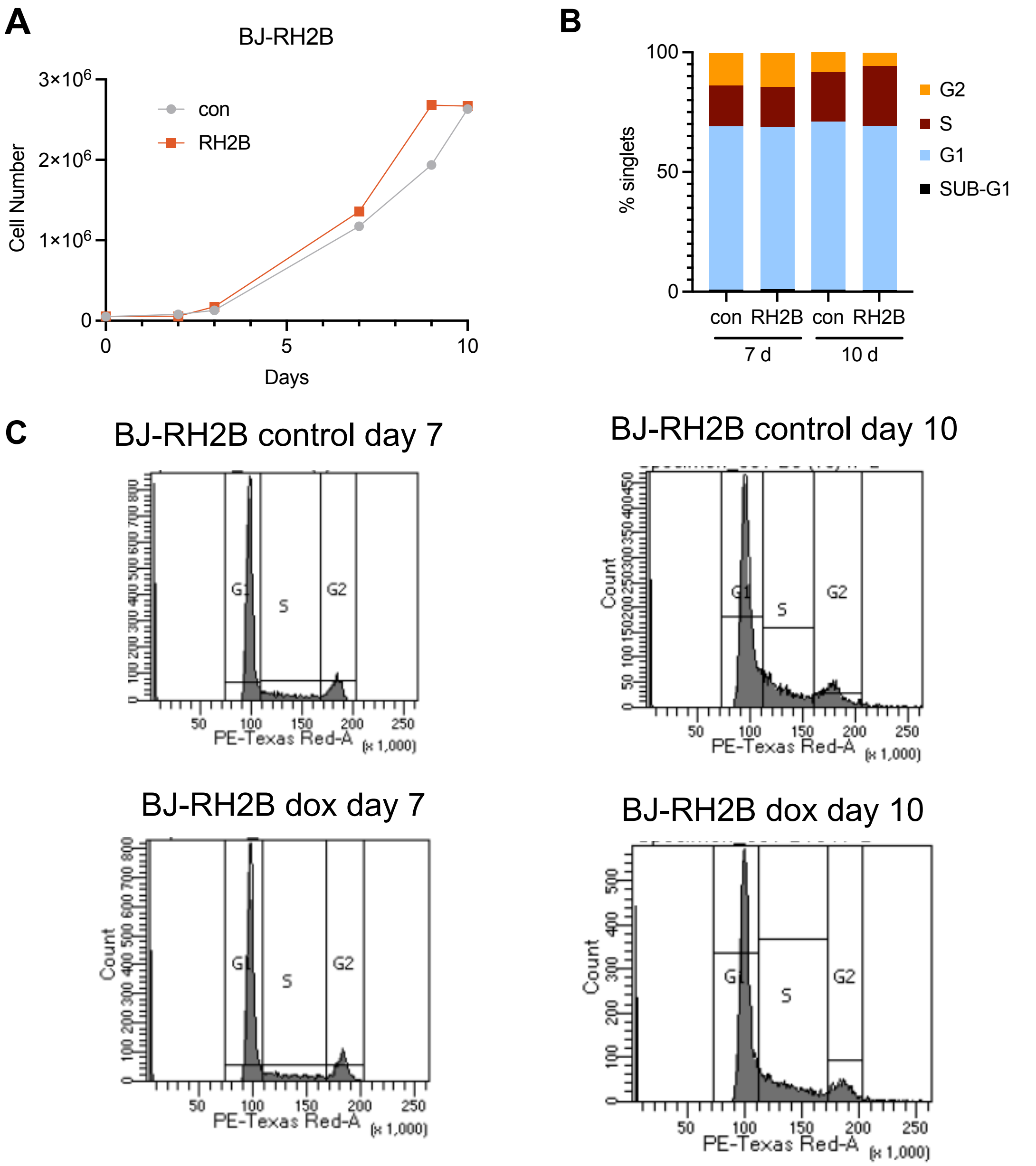


**Figure S5. Growth characteristics of cells overexpressing RNaseH2 subunits. (A)** Growth curve of BJ-RhH2B cells after RH2B induction or no DOX control (con), measured by cell number. N=1. **(B)** Quantification of cell cycle distribution in BJ-RH2B cells after 7 days or 10 days RH2B induction or no DOX control (con), based on measuring DNA content using propidium iodide staining and flow cytometry. N=1. **(C)** Flow cytometry gating strategy for quantification of cell cycle distribution as in B. Cells were stained with propidium iodide. The means and SEM (bars) of independent experiments are shown. Asterisks indicate p-values (student's t-test, ns: not significant).

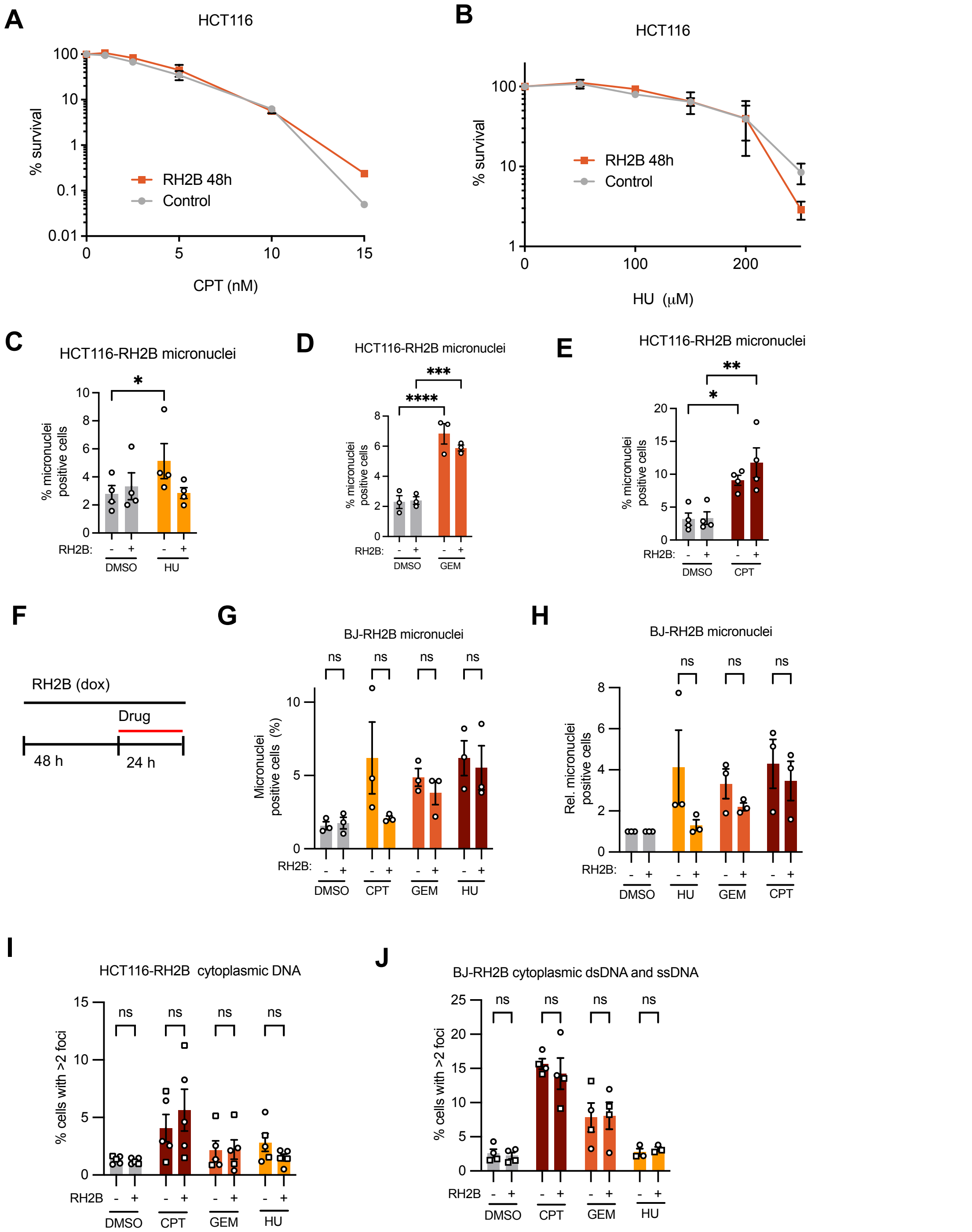


**Figure S6. Impact of RNaseH2 subunits on colony survival and genomic instability.**
**(A)** Colony survival of HCT116-RH2B cells after continuous treatment with CPT. N= 2. **(B)** Colony survival of HCT116-RH2B cells after continuous treatment with HU. N = 4. **(C)** Percentages of HCT116-RH2B cells with micronuclei +/- RH2B induction +/- 2mM HU. N=4. **(D)** Percentages of HCT116-RH2B cells with micronuclei +/- RH2B induction +/- 1 µM GEM. N=3. **(E)** Percentages of HCT116-RH2B cells with micronuclei +/- RH2B induction +/- 10 µM CPT. N=3. **(F)** Treatment schematic for micronucleus quantification in BJ-hTERT cells. **(G)** Percentages of BJ-hTERT-RH2B cells with micronuclei after 48 h RH2B induction and 24 h treatment with 10 µM CPT, 1 µM GEM, 200 µM HU or DMSO, and release from drug for 24 h. N=3. **(H)** Fold increase in BJ-hTERT-RH2B cells with micronuclei after 48 h RH2B induction and 24

h treatment with 10 µM CPT, 1 µM GEM, 200 µM HU or DMSO, and release from drug for 24 h. N=3. **(I)** Percentages of HCT116-RH2B cells with cytoplasmic ssDNA or dsDNA staining +/- 48 h RH2B induction and 24 h treatment with 1 mM CPT, 25 nM GEM, 200 mM HU or DMSO. Circles: dsDNA antibody, squares: ssDNA antibody. N=5. **(J)** Percentages of BJ-hTERT-RH2B cells with cytoplasmic ssDNA or dsDNA staining +/- 48 h RH2B induction and 24 h treatment with 1 mM CPT, 25 nM GEM, 200 mM HU or DMSO. Circles: dsDNA antibody, squares: ssDNA antibody. N=4 (CPT and GEM), N=3 (HU).

The means and SEM (bars) of independent experiments are shown. Asterisks indicate p-values (ANOVA, ns: not significant, * $p < 0.05$, ** $p < 0.01$, *** $p < 0.001$, **** $p < 0.0001$).

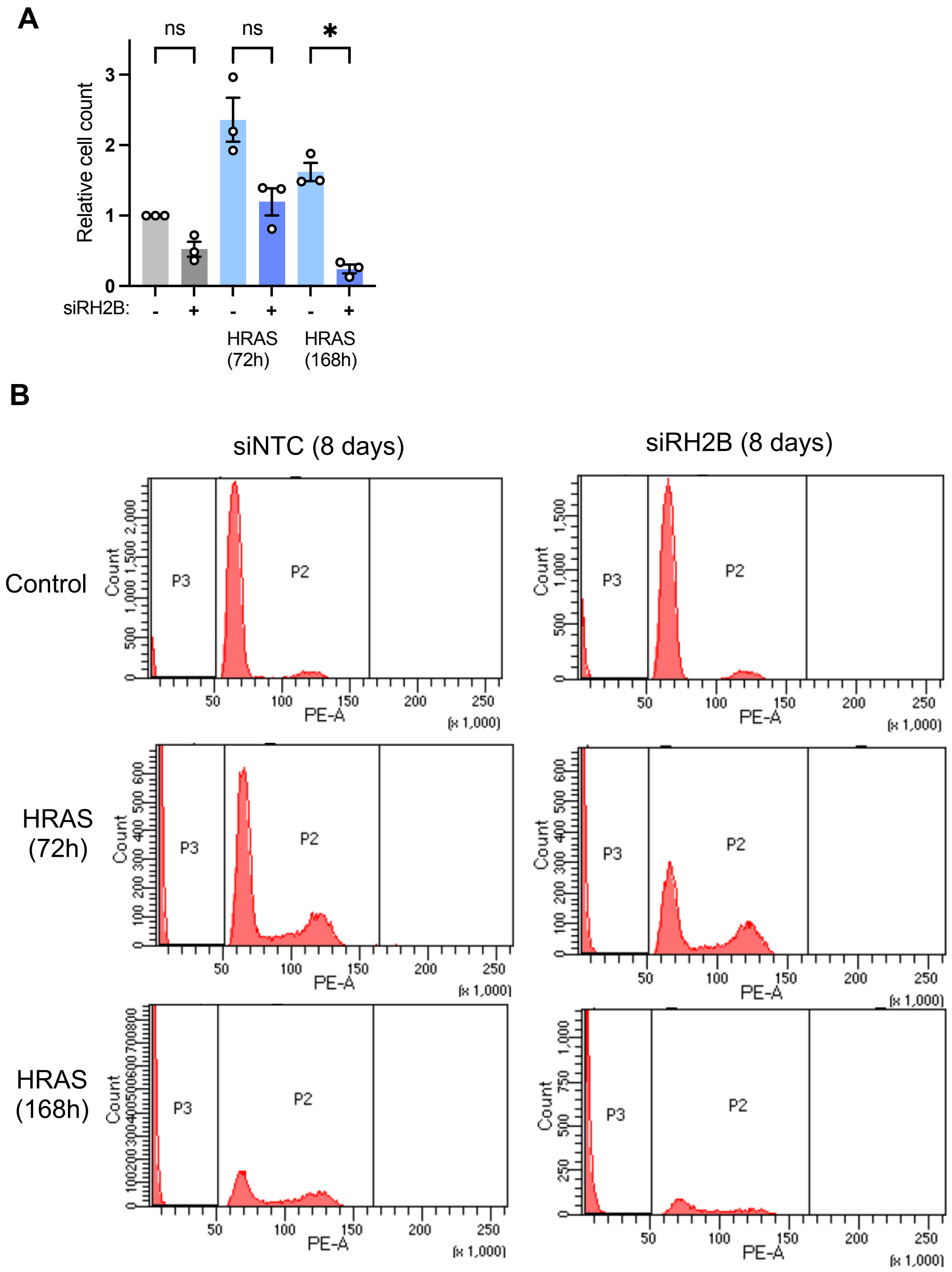


**Figure S7. RNaseH2B depletion exacerbates growth arrest and cell death in presence of oncogenic $HRAS^{G12V}$.**
**(A)** Relative proliferation after 8 days transfection with non-targeting control or RH2B siRNA and $HRAS^{G12V}$ induction for the times indicated. Cell counts were normalised to non-targeting control without $HRAS^{G12V}$ induction. N = 3.
**(B)** Flow cytometry gating strategy for quantification of sub-G1 phase populations. Cells were stained with propidium iodide. The means and SEM (bars) of independent experiments are shown. Asterisks indicate p-values (ANOVA, ns: not significant, * $p < 0.05$).
